# Runx3 prevents spontaneous colitis by directing differentiation of anti-inflammatory mononuclear phagocytes

**DOI:** 10.1101/742650

**Authors:** Shay Hantisteanu, Joseph Dicken, Varda Negreanu, Dalia Goldenberg, Ori Brenner, Dena Leshkowitz, Joseph Lotem, Ditsa Levanon, Yoram Groner

## Abstract

RUNX3 is one of three mammalian Runt-domain transcription factors (TFs) that regulate gene expression in several types of immune cells. Runx3-deficiency in mice is associated with a multitude of defects in the adaptive and innate immunity systems, including the development of early onset colitis. Our study reveals that conditional deletion of Runx3 specifically in mononuclear phagocytes (MNP) (MNP^Runx3−/−^) but not in T cells, recapitulates the early onset spontaneous colitis seen in Runx3^−/−^ mice.

We show that Runx3 is expressed in colonic MNP, including resident macrophages (RM) and the dendritic cell cDC2 subsets and its loss results in impaired differentiation/maturation of both cell types. At the transcriptome level, loss of Runx3 in RM and cDC2 was associated with upregulation of pro-inflammatory genes similar to those in the early onset IBD murine model of RM^Il10r−/−^. The impaired RM maturation in the absence of Runx3 was associated with a marked decrease in expression of anti-inflammatory and TGFβ-regulated genes. Similarly, the decreased expression of β-catenin signaling associated genes in Runx3-deficient cDC2 indicates their impaired differentiation/maturation. Analysis of ChIP-seq data suggests that in both MNP cell types a significant fraction of these differentially expressed genes are high confidence Runx3 directly regulated genes. Interestingly, several of these putative Runx3 target genes harbor SNPs associated with IBD susceptibility in humans. Remarkably, the impaired maturation and pro-inflammatory phenotype of MNP lacking Runx3 was associated with a substantial reduction in the prevalence of colonic lamina propria Foxp3^+^ regulatory T cells and an increase in IFNγ-producing CD4^+^ T cells, underscoring Runx3 critical role in establishing tolerogenic MNP.

Together, these data emphasize the dual role of Runx3 in colonic MNP, as a transcriptional repressor of pro-inflammatory genes and an activator of maturation-associated genes including anti-inflammatory genes. Our study highlights the significance of the current MNP^Runx3−/−^ model for understanding of human MNP-associated colitis. It provides new insights into the crucial involvement of Runx3 in intestinal immune tolerance by regulating colonic MNP maturation through TGFβR signaling and anti-inflammatory functions by Il10R signaling, befitting the identification of RUNX3 as a genome-wide associated risk gene for various immune-related diseases in humans including gastrointestinal tract diseases such as celiac and Crohn’s disease.

## INTRODUCTION

RUNX3 is one of the three mammalian Runt-domain transcription factors (TFs) that are key gene expression regulators in development (Levanon and Groner, 2004; Lotem et al., 2017). *Runx3* was originally cloned based on its similarity to *Runx1* (Levanon et al., 1994) and subsequently localized on human and mouse chromosomes 1 and 4, respectively (Avraham et al., 1995; Levanon et al., 1994). *Runx3*^−/−^ mice phenotypes reflect its expression pattern and essential requirement to the proper function of important organs (Ebihara et al., 2017; Lotem et al., 2017). Among these, absence of Runx3 is associated with a multitude of defects in the adaptive and innate immunity system including: defective proliferation and differentiation of activated cytotoxic CD8^+^ T cells (Cruz-Guilloty et al., 2009; Lotem et al., 2013; Taniuchi et al., 2002; Woolf et al., 2003); impaired induction of helper Th1 cells (Djuretic et al., 2007) and activation of natural killer (NK) cells (Levanon et al., 2014); impaired development of intestinal innate lymphoid cells type 3 (ILC3) (Ebihara et al., 2015); and lack of dendritic epithelial T cells (Woolf et al., 2007). We have also reported that Runx3 plays a pivotal role in mononuclear phagocytes (MNP) homeostasis. For example, Runx3 facilitates specification of splenic CD11b^+^ dendritic cells (DC) (Dicken et al., 2013) and is required for development of TGFβ-dependent skin Langerhans cells (Fainaru et al., 2004). Runx3^−/−^ bone marrow derived DC (BMDC) are hyper-activated due to deregulation of TGF-β-mediated maturation. Immune deficiencies in Runx3^−/−^ mice were also reported to cause lung inflammation associated with accumulation of hyper-activated DCs, leading to development of major hallmarks of asthma (Fainaru et al., 2005; Fainaru et al., 2004). Finally, Runx3^−/−^ mice spontaneously develop colitis at 4-6 weeks of age (Brenner et al., 2004). This disease is known to be closely associated with an impaired immune system. To this extent, it is of interest that human genome wide association studies have linked RUNX3 to various immune-related diseases including asthma, ankylosing spondylitis, psoriasis, psoriatic arthritis, atopic dermatitis and gastrointestinal tract (GIT) diseases such as celiac and Crohn’s disease (Lotem et al., 2017).

Expression of Runx3 in the GIT of WT mice is confined to leukocytes and not detected in GIT epithelium, indicating that cell autonomous expression of Runx3 in leukocytes is involved in GIT homeostasis (Levanon et al., 2011). In view of the defects in different innate and adaptive immune cell types in Runx3^−/−^ mice and the association of RUNX3 with immune related human diseases (Lotem et al., 2017), it became interesting to determine which of the Runx3^−/−^ immune cell types is directly involved in colitis development. Previously, it was shown that conditional deletion of Runx3 in NK and ILCs by crossing *Runx3^fl/fl^* on to *Nkp46-Cre* mice did not result in spontaneous colitis, although these mice did show a stronger intestinal damage following infection with C. *Rodentium* (Ebihara et al., 2015). Additionally, we have shown that *Runx3^−/−^* mice are highly resistant to inflammation-dependent skin chemical carcinogenesis and this resistance was fully recapitulated in Runx3 conditional knockout mice in which Runx3 was deleted in both DC and T cells, but not in epithelial cells (Bauer et al., 2014). Here we show that conditional deletion of Runx3 specifically in MNP, but not in T cells, recapitulates the spontaneous colitis seen in *Runx3^−/−^* mice. Specifically, Runx3 function in MNP is crucial for intestinal immune tolerance by regulating their proper maturation and anti-inflammatory functions.

## MATERIALS AND METHODS

### Mice

Mice lacking Runx3 specifically in MNP were generated by crossing *Runx3*^*fl/fl*^ mice (Levanon et al., 2011) onto *CD11c-Cre* (Caton et al., 2007) or *Cx3cr1c-Cre* mice (Yona et al., 2013), giving rise to *Runx3*^*fl/fl*^*/CD11c:Cre* (hereafter called *Runx3*^**Δ**^) or *Runx3*^*fl/fl*^*/Cx3cr1:Cre* (*Cx3cr1-Runx3*^*Δ*^) mice, respectively. *Runx3*^**Δ**^ mice were crossed onto *Cx3cr1-GFP* mice (Jung et al., 2000) to obtain *Runx3*^**Δ**^ Cx3cr1-GFP mice. All animals were on the C57Bl/6 background. Mice lacking Runx3 specifically in T-cells were generated by crossing *Runx3*^*fl/fl*^ mice onto *Lck-Cre* mice (Takahama et al., 1998) (*Runx3*^*fl/fl*^*/Lck:Cre)*. The *Runx3*^*P1-AFP/P2-GFP*^ knock-in (KI) named here Runx3-GFP and *Runx3*^*−/−*^ mice were previously described (Levanon et al., 2011). C57Bl/6 Ly5.2 mice were purchased from Harlan Laboratories (Rehovot). C57Bl/6 Ly5.1 mice were bred in the Weizmann animal facility. To determine the ability of Runx3^−/−^ leukocytes to transfer colitis, we adoptively transferred intravenously 3×10^6^ E13.5 fetal liver (FL) cells from WT or *Runx3*^*−/−*^ C57Bl/6 mice into lethally irradiated (800R and 400R, 4 h apart) C57Bl/6 mice and colitis pathology was determined 2 months post transfer. For generation of BM chimeras, C57Bl/6 CD45.1 mice were lethally irradiated (1050R) and reconstituted by intravenous injection of a 1:1 mixture of WT C57Bl/6 CD45.1 and CD11c-Runx3^Δ^ CD45.2 BM cells. Mice were analyzed 11-15 weeks post BM transfer.

Animals were maintained under SPF conditions and handled in accordance with the protocol approved by the Institutional Animal Care and Use Committee (IACUC) of the Weizmann Institute of Science (Permit #: 09750119-4).

Genotyping primers; Floxed: F5’-CCCACCCATCCAACAGTTCC, R5’-GAGACCACAGAGGACTTGTA. CRE: F5’-AACATGCTTCATCGTCGG, R5’-TTCGGATCATCAGCTACACC. Cx3cr1-GFP mice WT or GFP allele: F5’-TTCACGTTCGGTCTGGTGGG, R5’-GGTTCCTAGTGGAGCTAGGG and F5’-GATCACTCTCGGCATGGACG, R5’-GGTTCCTAGTGGAGCTAGGG, respectively.

### Isolation and analysis of colonic lamina propria (LP) cells

Isolation of colonic LP cells was performed as previously described with some modifications (Varol et al., 2009). Briefly, extra-intestinal fat tissue and blood vessels were carefully removed and colons were then flushed of their luminal content with cold PBS. The cecum was opened longitudinally, and cut into 0.5 cm pieces. Epithelial cells and mucus were removed by 40 min incubation at 37°C with shaking at 250 rpm in Hank’s balanced salt solution containing 5% fetal bovine serum (FBS) and 2 mM EDTA. Colon pieces were then digested by 40 min incubation at 37°C with shaking in RPMI-1640 containing 5% FBS, 1 mg/ml Collagenase II (Worthington, US) and 0.1 mg/ml DNase I (Roche, US). The digested cell suspension was then washed with PBS and passed through 100 μm cell strainer. For analysis of blood monocytes, samples were resuspended in ACK erythrocyte-lysis buffer (0.15M NH4Cl, 0.1M KHCO3 and 1mM EDTA in PBS). For intracellular staining, a Foxp3 buffer set (Invitrogen, US) was used according to manufacture instructions. To induce T-cell activation for determination of intracellular IFN-γ, LP cells were incubated for 5 h with 20 ng/ml PMA and 1 μg/ml ionomycin in the presence of 10 μg/ml Berfeldin A (Sigma, IL). For FACS analysis, single cell suspensions were stained with the following antibodies (Abs); Epcam G8.8, CD16/32 clone 93, CD45 30-F11, MHCII M5/114.15.2, CD11c N418, Ly6c HK1.4, CD11b M1/70, CD103 2E7, F4/80 BM8, CD64 X54-5/7.1, CD115 AFS98, CD43 eBioR2/60, Foxp3 MF-14, CD25 PC61, CD45RB C363-16A, CD4 RM4-5, Clec12a 5D3, PD-L2 TY25, CD24a M1/69, T-bet 4B10, IFN-γ XMG1.2, CD45.1 A20 and CD45.2 104. All Abs were purchased from BioLegend, US or eBiosciences, US unless indicated otherwise. LSRII flow cytometer (BD Biosciences, US) with FACSDiva Version 6.2 software (BD Biosciences, US) was used and further data analysis was conducted using FlowJo Version 9.7.6 software (TreeStar, BD US). FACSAria flow cytometer (BD Biosciences, US) was used for cell sorting and forward scatter height versus forward scatter width appearance was used to exclude doublets.

### Immunofluorescence (IF) and Immunohistochemistry (IHC)

Colon was removed and frozen in liquid nitrogen. Cryo-sections of 12-14 μm were prepared on glass slides, fixed for 3 min in acetone at −20°C and air dried for 20 min. Slides were blocked with PBS containing 20% horse serum and stained O.N at RT with rabbit anti-MHCII-biotin Ab followed by incubation with SA-cy3 Ab. MHCII staining and endogenous Cx3cr1-GFP signal was analyzed by fluorescent microscope. In addition, 4 μm of paraffin serial sections of colon and stomach were prepared and stained with hematoxylin-eosin (H&E) for histopathology evaluation as described (Brenner et al., 2004).

### RNA extraction and microarray gene expression analysis

Total RNA was extracted from sorted cecum MNP using the RNeasy Mini Kit (QIAGEN, US). In each experiment sorted cells were pooled from 3-4 mice. BioAnalyzer 2100 (Agilent Technologies, US) was used to determine RNA quality. RNA from each sample was labeled and hybridized to Affymetrix mouse exon ST 1.0 microarrays according to manufacturer instructions. Microarrays were scanned using GeneChip scanner 3000 7G. Statistical analysis was performed using the Partek® Genomics Suite (Partek Inc., US) software. CEL files (raw expression measurements) were imported to Partek GS and the data was preprocessed and normalized using the RMA (Robust Multichip Average) algorithm (Irizarry et al., 2003) with GC correction. To identify differentially expressed genes (DEGs), One-Way ANOVA was applied. DEGs lists were created by filtering the genes based on absolute fold change ≥ 1.5, p ≤ 0.05. Log_2_ gene intensities were used for volcano scatter plots (Partek). Ingenuity software was applied for GO analysis. See below GEO accession number.

### Chromatin Immunoprecipitation Sequencing (ChIP-seq) data acquisition and analysis

Two biological replicate ChIP-seq experiments were conducted for detection of Runx3-bound genomic regions. 30×10^6^ cells of the D1 DC cell line were fixed in 1% formaldehyde and sonicated to yield DNA fragments of ~300 bp according to standard procedures previously described (Pencovich et al., 2011). For immunoprecipitation (IP), 40 μl of anti-Runx3 Ab were added to 15 ml of diluted fragmented chromatin and incubated overnight at 4°C; Rabbit pre-immune serum was used as control. DNA was purified using QIAquick spin columns (QIAGEN, US).

For ChIP-seq analysis, Illumina sequencing of short reads was performed in Illumina Genome Analyzer. For data analysis, Runx3 IP and control IP extracted sequences were aligned uniquely to the mouse genome (mm9) using bowtie (Langmead et al., 2009). Bound regions were detected using MACS2 (Feng et al., 2012). Runx3 bound peaks and coverage data (bigWig files) were uploaded to the UCSC genome browser. GREAT algorithm (version 3.0.0) (McLean et al., 2010) was applied to determine genes corresponding to Runx3-bound peaks in D1 cells and splenic CD4^+^ DCs (Dicken et al., 2013) and to H3K4me1, H3K27Ac and ATAC-seq peaks in colonic RM (Lavin et al., 2014). Cistrome CEAS (Liu et al., 2011) was used to compute enrichment of genomic sequences. Discovery of TF binding sites was conducted using Genomatix (https://www.genomatix.de). All microarray and ChIP-seq data are available in the GEO public database under the SuperSeries accession number GSE136067.

### RT–qPCR and protein analysis

Total RNA was reverse-transcribed using miScript reverse transcription kit (QIAGEN) according to manufacturer’s instructions. Quantitation of cDNAs was performed by applying sequence-specific primers (see below), miScript SYBR Green PCR kit (QIAGEN) and using Roche LC480 *LightCycler*. Target transcript quantification was calculated relative to *Hprt* mRNA. Standard errors were calculated using REST (Pfaffl et al., 2002).

qPCR Primers:

Runx3 ex3-4 F: 5’-GCCGGCAATGATGAGAACT

Runx3 ex3-4 R: 5’-CACTTGGGTAGGGTTGGT

Hprt-F: 5’-GTTGGATACAGGCCAGACTTTGTTG

Hprt-R: 5’-CCAGTTTCACTAATGACACAAACG

Hpgds-F: 5’-GGAAGAGCCGAAATTATTCGCT

Hpgds-R: 5’-ACCACTGCATCAGCTTGACAT

Ccl24-F: 5’-ACCGAGTGGTTAGCTACCAGTTG

Ccl24-R: 5’-TGGTGATGAAGATGACCCCTG

Fcrls-F: 5’-ACAGGATCTAAGTGGCTGAATGT

Fcrls-R: 5’-CTGGGTCGTTGCCCTATCTG

Cxcl9-F: 5’-TCCTTTTGGGCATCATCTTCC

Cxcl9-R: 5’-TTTGTAGTGGATCGTGCCTCG

Ms4a14-F: 5’-ACCAACAGACCAGCAGTCAGAAGA

Ms4a14-R: 5’-TTGGATGAGCCTGAGCAAGGTGTA

Pd-l2 –F: 5’-CTGCCGATACTGAACCTGAGC

Pd-l2 –R: 5’-GCGGTCAAAATCGCACTCC

For protein analysis, cell proteins were extracted with Radioimmunoprecipitation assay (RIPA) buffer containing protease inhibitors and analyzed by western blotting using either in-house anti-Runx3 or anti-Runx1 Ab. GAPDH was used as an internal loading control.

### Statistical analysis

Statistical significance was determined with unpaired, two tailed Student’s t-test.

## RESULTS

### Loss of Runx3 in non-T cells leukocytes induces colitis

To establish whether Runx3^−/−^ leukocytes are involved in colitis development, we performed a transfer experiment. Because *Runx3*^*-/-*^ mice on C57Bl/6 background are not viable after birth, we adoptively transferred E13.5 FL cells from either *Runx3*^*-/-*^ or WT mice into lethally irradiated C57Bl/6 mice. Histopathological analysis performed at 2 months post transfer revealed that recipients of Runx3^−/−^ but not WT FL cells developed inflammatory bowel disease (Figure 1A, B), in agreement with the results of Sugai et al (Sugai et al., 2011). To determine whether this phenotype can be attributed to T cells, we crossed *Runxtf3*^*fl/fl*^ mice on to *Lck-Cre* mice to obtain T-cell specific Runx3 conditional knockout (cKO) mice. Unlike the spontaneous colitis seen in *Runx3*^*-/-*^ mice, *Lck-Runx3*^*Δ*^ mice did not develop spontaneous colitis (Figure 1C, D). As indicated earlier, colitis also did not occur in *NKp46-Runx3*^*Δ*^ mice (Ebihara et al., 2015). These results indicate that Runx3 function in T, NK and ILC is not essential to maintain GIT homeostasis and suggest that cell autonomous Runx3 function in other immune cell types, possibly MNP, is involved.

**Figure 1.**
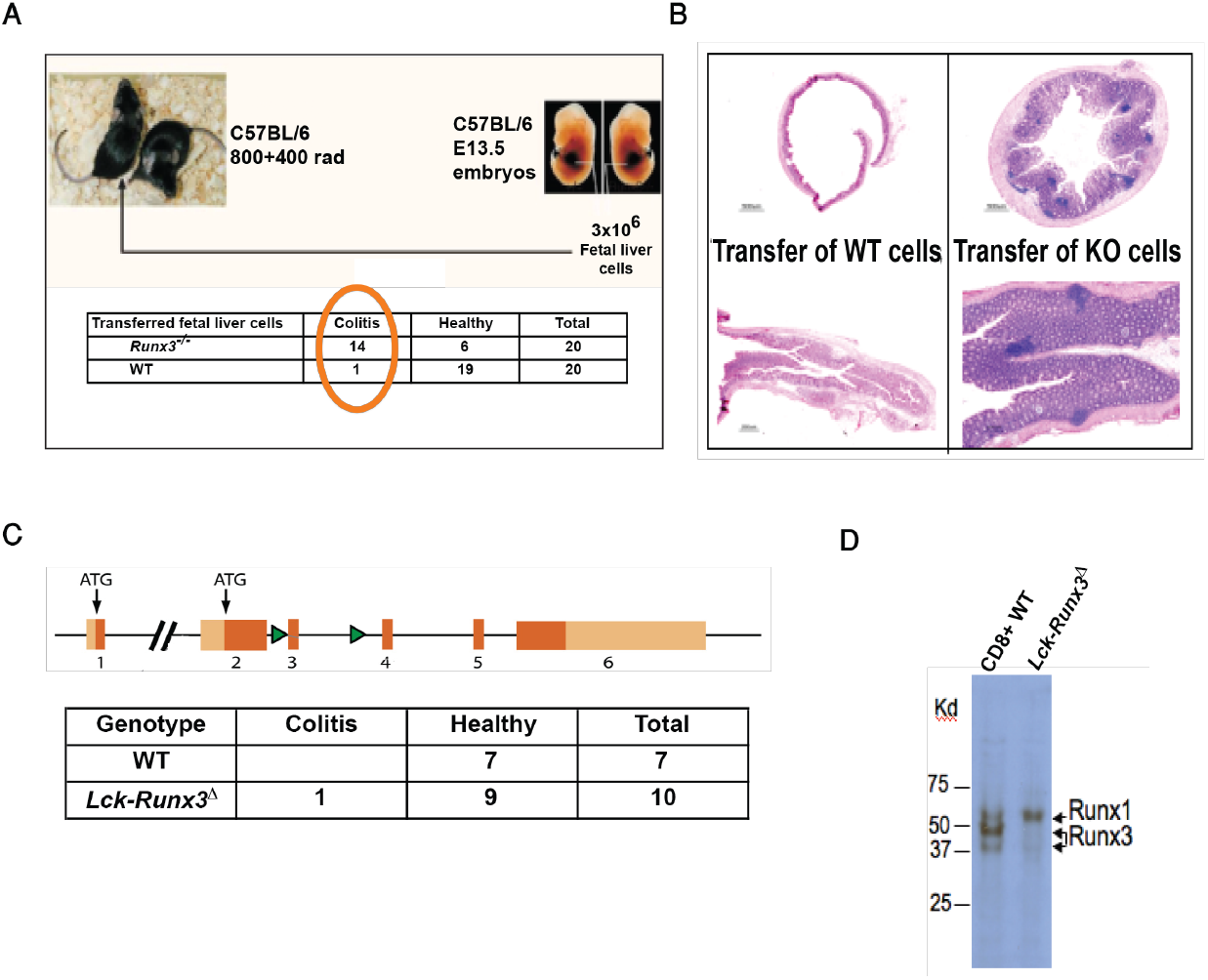
Loss of Runx3 in non-T cells leukocytes induces colitis. **A-B**, Adoptive transfer of Runx3^*−/−*^ but not WT FL cells induces colitis in lethally irradiated recipient mice. **A**, Number of recipient mice with colitis. **B**, Histological sections of colon stained with H&E. **C**, T cell-specific ablation of Runx3 *(Lck-Runx3*^*Δ*^) does not induce colitis, shown in the table. Upper, schematic representation of *Runx3* gene locus targeted for conditional inactivation. Orange dark boxes represent coding exons, light orange boxes are UTRs, exons are marked by numbers. The green arrowheads represent lox-P sites that flank exon 3, one of the exons comprising the RUNX TF DNA binding domain. **D**, Western blot of CD8^+^ T cell extract from WT and *Lck-Runx3*^**Δ**^ mice. Note the efficient deletion of Runx3 and up-regulation of Runx1 in *Lck-Runx3*^**Δ**^ cells.

### Mice lacking Runx3 in colonic MNP develop early onset spontaneous colitis

MNP are known for their role in maintaining GIT homeostasis (Joeris et al., 2017). In order to determine whether Runx3 expression in MNP plays role in GIT immune tolerance, we generated MNP-specific Runx3-cKO mice by crossing *Runx3*^*fl/fl*^ mice onto two different Cre transgenic strains, *CD11c-Cre* and *Cx3cr1-Cre*, generating *Runx3*^**Δ**^ and *Cx3cr1-Runx3*^**Δ**^ mice, respectively. CD11c and Cx3cr1 are expressed in both DC and macrophages, but CD11c is expressed at a higher level in DC and Cx3cr1 at a higher level in macrophages. Strikingly, the majority of ~3 months old mice of both MNP-specific *Runx3*-cKO mouse models developed spontaneous mild colitis affecting the cecum and proximal colon (Figure 2A, B). Colitis in these *Runx3*-cKO mice displayed similar characteristics to those observed at an earlier age in *Runx3^−/−^* mice (Brenner et al., 2004), including accumulation of infiltrating leukocytes associated with pronounced mucosal hyperplasia and loss of differentiated mucous secreting goblet cells. *Runx3*^**Δ**^ and *Cx3cr1-Runx3*^**Δ**^ mice older than 7 months generally showed exacerbated colon inflammation and about 30% of the mice also developed gastropathy (Figure 2C), resembling the phenotype of *Runx3^−/−^* mice (Brenner et al., 2004). Of note, younger mice aged 6-8 weeks showed low prevalence of colitis with a minimal average disease score.

**Figure 2.**
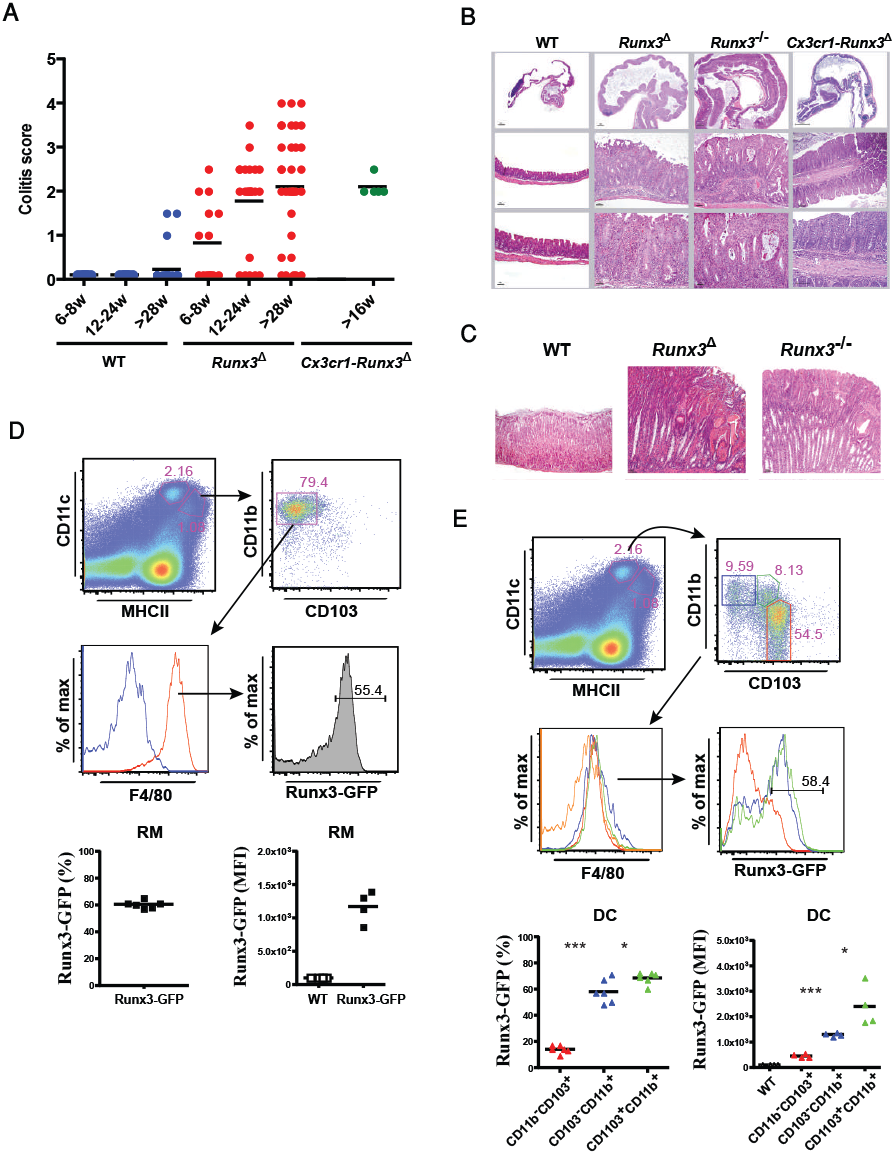
Colitis development in Runx3^*−/−*^ and in the two MNP-specific *Runx3-cKO* mice (*Runx3*^**Δ**^ and *Cx3cr1-Runx3*^**Δ**^). **A**, A graphical summary of postmortem colitis grade in *Runx3*^**Δ**^ and *Cx3cr1-Runx3*^**Δ**^ (right) relative to WT mice (left). Colitis score was scaled from 1 to 4 (1-minimal; 2-mild; 3-moderate; 4-severe). Each dot represents one mouse. (Note the complete absence of pathological signs of colitis in WT mice 24 weeks of age). **B**, H&E stained histological sections of colons from WT and the three Runx3-deficient mouse strains. **C**, H&E stained histological sections of stomach of two Runx3-deficient 1-year old mouse strains compared to WT. **D**, Flow cytometry analysis of *Runx3* expression in colonic RM of Runx3-GFP^+^ mice. The RM were identified by F4/80 expression (red line) and overly on F4/80^−^CD11b^+^ (blue line) (top and middle panels). Graphical summary of Runx3-GFP+ RM (bottom, left panel) and mean fluorescence intensity (MFI) relative to non-GFP (WT) control (bottom, right panel). **E**, Flow cytometry analysis of *Runx3* expression in colonic CD103^+^CD11b^−^ cDC1 (red), CD103^−^CD11b^+^ cDC2 (blue) and CD103^+^CD11b^+^ cDC2 (green) (top and middle panels). Orange line represents the F4/80 negative level. Graphical summary of Runx3-GFP+ prevalence among the DC subsets (bottom left) and Runx3-GFP^+^ MFI relative to non-GFP (WT) control (bottom right).

Since colitis developed in the MNP-specific *Runx3*-cKO mice, we determined which subtype of colonic LP MNP cells expresses *Runx3* by employing compound heterozygous mice (Runx3-GFP) (Dicken et al., 2013). Analysis revealed that within the LP MNP compartment *Runx3* is expressed in resident macrophages (RM), characterized as CD11c^int^CD11b^+^MHCII^+^F4/80^+^CD103^−^ (Figure 2D), and also at a low level in LP CD11c^lo^ monocytes (Figure S1A), but not in circulating monocytes (CD11b^+^CD115^+^CD43hiCD11c^+^Ly6c^−^ and CD11b^+^CD115^+^CD43loCD11c-Ly6c^+^) (Figure S1B). These results are in agreement with the identification of Runx3 as an intestinal RM specific TF (Lavin et al., 2014) and indicate that Runx3 expression accompanies the differentiation of monocytes to colonic RM.

In addition to RM, the MNP compartment contains three DC subsets commonly characterized as CD11c^high^MHCII^+^F4/80^low^. These subsets include CD103^+^CD11b^−^ termed conventional DC1 (cDC1) and two cDC2 subsets: CD103^−^CD11b^+^ and CD103^+^CD11b^+^ DC. We detected expression of *Runx3* in the majority of CD103^−^ CD11b^+^ and CD103^+^CD11b^+^ DC, while only a small cDC1s fraction expressed *Runx3*. Moreover, these Runx3-expressing cDC1s had a lower *Runx3* level relative to the two CD11b^+^ DC subsets (Figure 2E). Recent work studying the relationship between intestinal DC subsets suggested that CD103^+^CD11b^+^ DC develop from intermediate CD103^+^CD11b^+^ DC in a TGF-β-dependent manner (Bain et al., 2017). Accordingly, CD103^+^CD11b^+^ DC expressed higher *Runx3* level compared to CD103^+^CD11b^+^ DC (Figure 2E). Taken together, our results imply that Runx3^−/−^ CD11b^+^ MNPs, including both RM and CD11b^+^ DC, drive the spontaneous development of colitis. Nevertheless, the contribution of the small fraction of Runx3-expressing CD103^−^CD11b^+^ DC cannot be ruled out. Using *Runx3-P1*^*AFP/+*^ or *Runx3-P2*^*GFP/+*^ reporter mice established that in CD103^+^CD11b^+^ DC and in RM both promoters mediate *Runx3* expression, however, P1 or P2 were preferentially used in these cells, respectively (Figure S1C). Collectively, the results indicate that the known role of intestinal MNP in maintenance of GIT homeostasis (Joeris et al., 2017) is mediated, at least in part, by P1 and P2 driven *Runx3* expression.

### Loss of Runx3 in MNP causes an early imbalance of colonic MNP subsets

As loss of Runx3 in MNP triggers spontaneous colitis, we sought to determine whether alterations in the colonic MNP compartment in *Runx3*^**Δ**^ mice occur prior to the onset of colitis. Colon RM are derived from recruited circulating monocytes, which differentiate to RM in four stages (S) (SP1 to SP4) described as the “waterfall” differentiation pattern (Bain et al., 2013; Schridde et al., 2017). Under steady state conditions in WT mice, the mature anti-inflammatory SP3 and SP4 stages predominate (Figure 3A). In *Runx3*^**Δ**^ mice at 6-8 weeks of age there was a marked increase in the prevalence of Ly6c^+^ pro-inflammatory SP2 monocytes with a concomitant decrease in the mature SP3+SP4 fraction (Figure 3A). These results suggest that Runx3 function in MNP is required for the maturation of colonic monocytes into anti-inflammatory RM. The transition from SP3 to fully mature SP4 RM is characterized by increased expression of Cx3cr1 (Schridde et al., 2017). To evaluate the level of Cx3cr1 expression in Runx3^**Δ**^ versus WT RM, we generated *CD11c-Runx3*^**Δ**^*-Cx3cr1^GFP^* mice. Colonic sections showed more cells expressing Cx3cr1^GFP^ signal in *Runx3*^**Δ**^ compared to WT. The GFP signal in the *Runx3*^**Δ**^ colon sections was scattered uniformly in the LP, whereas in WT colon sections these cells were located relatively adjacent to the epithelium (Figure 3B). Moreover, GFP intensity analysis revealed a significant decrease in the number of Cx3cr1^GFP-hi^ SP4 cells and an increased number of the less mature Cx3cr1^GFP-int^ SP3 cells in the *Runx3*^**Δ**^ LP (Figure 3C, S2A). These results indicate that Runx3 is involved in the terminal maturation of RM from the SP3 to the anti-inflammatory SP4 stage.

**Figure 3.**
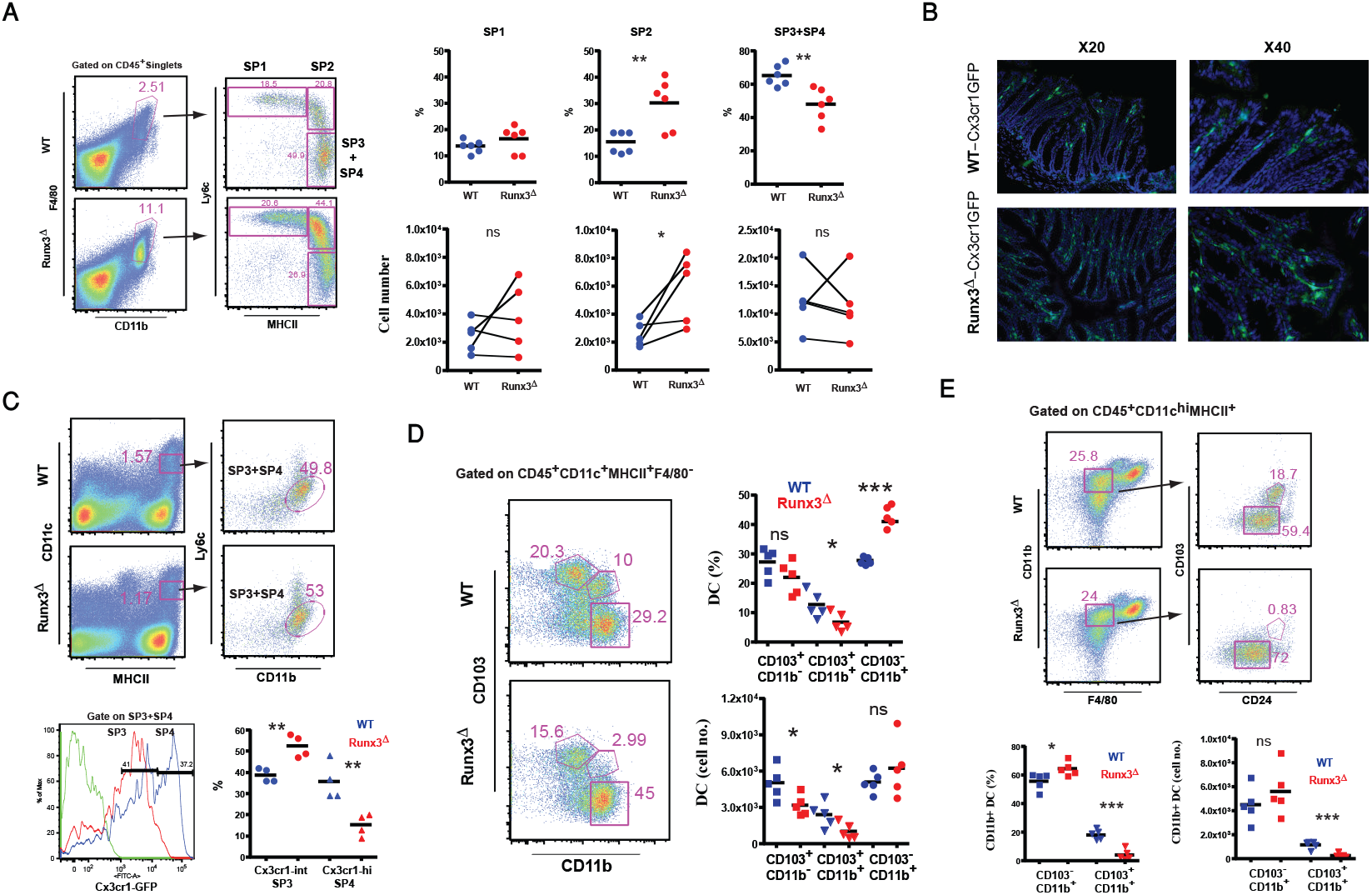
Loss of Runx3 in MNP causes an imbalance among colonic MNP subsets. **A**, Comparison of WT and *Runx3*^**Δ**^ colonic monocyte-macrophage “waterfall”. Representative flow cytometry gating on CD45^+^CD11b^+^F4/80^+^ cells demonstrating SP1, SP2 and SP3+SP4 cells (left). Graphical summary comparing the prevalence and cell numbers of WT and *Runx3*^***Δ***^ colon SP1, SP2 and SP3+SP4 waterfall cells (right). **B**, GFP signal in colon frozen section of 6-8 weeks old *Runx3*^**Δ**^-*Cx3cr1*^*GFP/+*^ mice compared to WT-Cx3cr1^GFP/+^ littermates. **C**, Analysis of Cx3cr1-GFP expression level in WT RM relative to *Runx3*^**Δ**^ colonic RM. Top, representative flow cytometry gating on WT and *Runx3*^**Δ**^ CD45^+^CD11c^int^MHCn^+^CDnb^+^Ly6c^−^ RM. Bottom, overlay histogram (left) of Cx3cr1-GFP expression level in WT (blue) and *Runx3*^**Δ**^ (red) RM and graphical summary (right) of the frequency of Cx3cr1^Hi^ and Cx3cr1^Int^ macrophages in WT and Runx3^A^. **D**, DC subsets in WT and *Runx3*^*Δ*^ colonic LP of 7-8 weeks old mice. Left, a representative flow cytometry of the CD103^+^CD11b^−^, CD103^+^CD11b^+^ and CD103^−^CD11b^+^ DC subsets. Right, graphical summary depicting the prevalence and cell numbers of the three colonic LP DC subsets. **E**, Representative flow cytometry gating on CD45^+^CD11c^hi^MHCII^+^CD11b^+^F4/80^lo^cells comparing CD24a expression between *Runx3*^**Δ**^and WT at 8 weeks of age (upper). Graphical summary comparing the prevalence and cell numbers of WT and Runx3^**Δ**^ CD103^+^CD11b^+^CD24a^+^ and CD103^−^CD11b^+^CD24a^+^ DC (lower).

Analysis of 7-8 weeks old *Runx3*^**Δ**^ mice revealed that among the 3 DC subsets, the CD103^+^CD11b^+^ and CD103^−^CD11b^+^ DC subsets were decreased and increased, respectively, compared to WT littermates, while the CD103^+^CD11b^−^ DC subset was only slightly reduced. Furthermore, quantification of DC subsets displayed a significant reduction in the number of CD103^+^CD11b^+^ DC in the *Runx3*^**Δ**^ mice (Figure 3D). Employing CD24a, an additional marker of CD103^+^CD11b^+^ DC, detected even more substantial reduction in prevalence and number of CD103^+^CD11b^+^ DC (Figure 3E). To determine whether the above documented decrease in abundance of CD103^+^CD11b^+^ DC in *Runx3*^**Δ**^ compared to WT mice occurs earlier than at 7-8 weeks, we examined 5-6 weeks old mice. Interestingly, at this younger age the *Runx3*^**Δ**^ mice showed no difference in DC subsets distribution and cell number compared to WT (Figure S2B), corresponding with the pathology data in Figure 2A. However, *Runx3*^*Δ*^ CD103^+^CD11b^+^ DC displayed reduced level of both CD24a and CD103 (Figure S2B and S2C). Thus, these results suggest that Runx3 loss in *Runx3*^**Δ**^ mice causes an imbalance among colonic MNP subsets and disturbs development of CD103^+^CD11b^+^ DC, prior to the onset of significant colitis. These changes are associated with the loss of anti-inflammatory and gain of pro-inflammatory RM and DC populations, reflected by the increased prevalence of colonic pro-inflammatory SP2 monocytes and CD103^−^CD11b^+^ DC and decrease of the anti-inflammatory SP4 RM and CD103^+^CD11b^+^ DC. Overall, the findings support a cell-autonomous Runx3 function in colonic CD11b^+^ MNP, emphasizing their importance to intestinal homeostasis. Accordingly, loss of Runx3 in MNP causes an early impairment of colonic CD11b^+^ MNP, marking it as an early onset inflammatory sign.

### Transfer of WT BM overcomes Runx3^Δ^ BM-mediated colitis onset

Competitive BM-repopulation assays were conducted to address whether Runx3-sufficient MNP regulate an anti-inflammatory response or whether in the presence of WT MNP, Runx3^**Δ**^ MNP will still dictate a pro-inflammatory response. Reconstituted mice colons were examined histopathlogically for colitis signs and by flow cytometry to assess the replenishment of colon MNP compartment. A 1:1 mixture of Runx3^**Δ**^ BM CD45.2 and WT BM CD45.1 cells was transferred into WT CD45.1 recipient mice. Eleven to fifteen weeks following transfer, colons of experimental mixed chimeric (Runx3^**Δ**^/WT→WT) mice were compared to those of Runx3^**Δ**^→WT and WT→WT reconstituted mice. There were no differences in weight gain between the different recipient mice groups during the course of the transfer experiment (Figure S3A). As could have been predicted, colons of Runx3^**Δ**^ BM reconstituted mice displayed colitis development (Figure 4A). Intriguingly, however, the mixed chimeric BM reconstituted mice did not develop colitis (Figure 4A). This finding underscores the ability of Runx3-sufficient MNP to dictate a mucosal anti-inflammatory environment that overrides the pro-inflammatory state of Runx3^**Δ**^ MNP.

**Figure 4.**
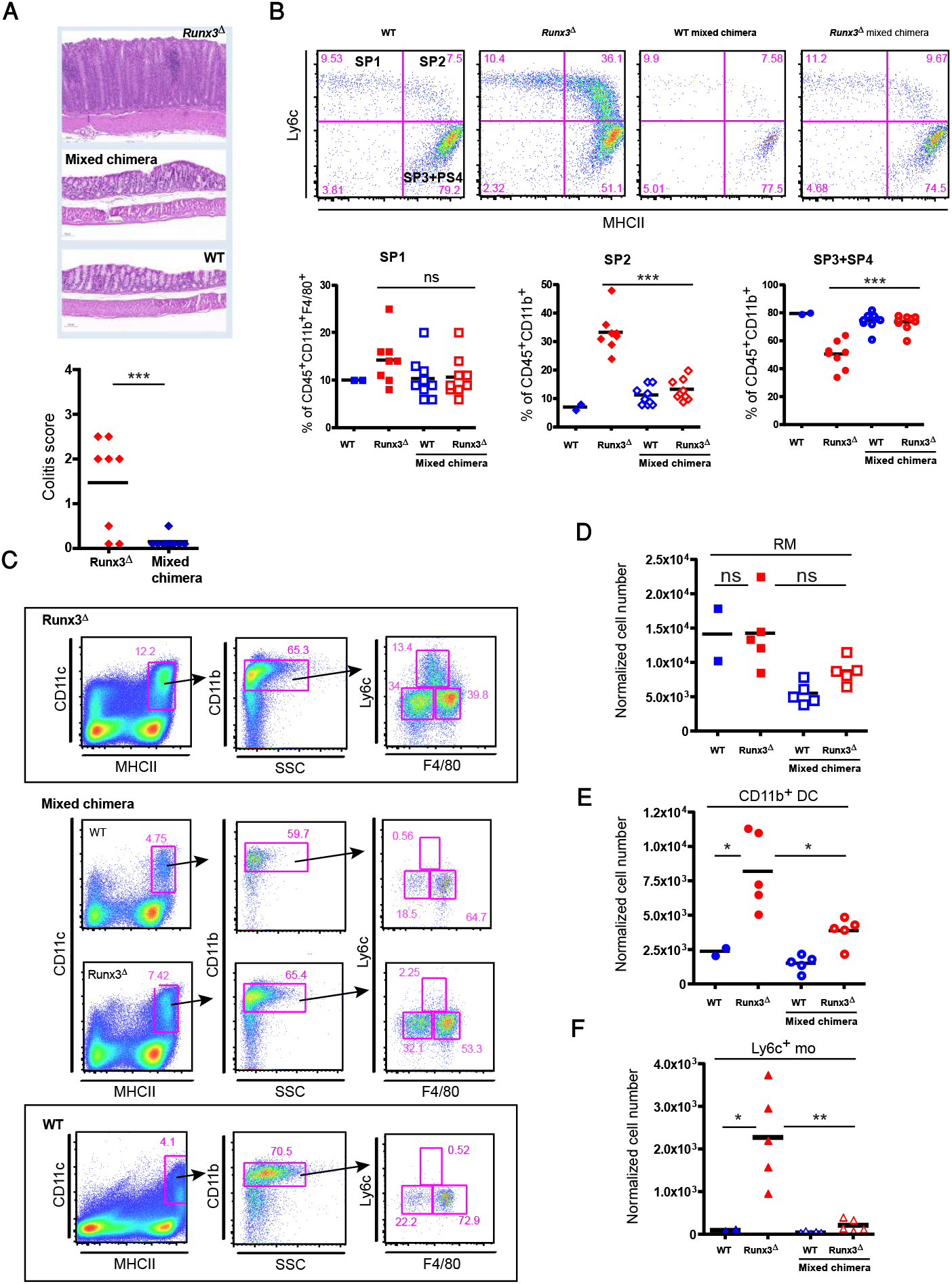
Colitis and colonic MNP subsets in lethally irradiated mice reconstituted with WT, *Runx3*^*Δ*^ or a mixture of *WT/Runx3*^*Δ*^ BM cells. **A**, Representative H&E stained colon section of Runx3^**Δ**^ BM recipient mice [Runx3^**Δ**^ (CD45.2) →WT(CD45.1)], mixed chimera [Runx3^**Δ**^ (CD45.2)/WT (CD45.1)→ WT(CD45.1)] and WTBM recipient mice [WT (CD45.1)®WT (CD45.1)] (top). Colitis score was scaled from one to four (1-minimal; 2-mild; 3-moderate; 4-severe). *** p<0.001 (bottom). **B**, Comparison of colonic LP SP1 to SP4 waterfall between WT, Runx3^**Δ**^ and mixed chimeras (WT and Runx3^**Δ**^) BM recipient mice. Representative waterfall from each group (top) and summary of SP1, SP2 and SP3+SP4 abundance in the three groups (bottom) are shown. *** p<0.001. C, Representative flow cytometry profile of colonic LP MNP subsets in *CD11c-Runx3*^**Δ**^ BM recipient mouse (top), mixed chimeric mouse (middle) and WTBM recipient mouse (bottom). **D**, Graphical summary of colonic LP RM normalized cell number in reconstituted BM mixed chimera compared to Runx3^**Δ**^ BM reconstituted mice and WTBM reconstituted mice. **E**, Graphical summary of colonic LP CD 11b^+^ DC normalized cell number in reconstituted BM mixed chimera compared to Runx3^**Δ**^ BM reconstituted mice and WTBM reconstituted mice. * p<0.05. **F**, Graphical summary of colonic LP Ly6c^+^ monocytes normalized cell number in reconstituted BM mixed chimera compared to Runx3^**Δ**^ BM recipient and WTBM recipient mice. * p<0.05, ** p<0.01.

Analysis of colon RM waterfall in Runx3^**Δ**^ BM recipients revealed a marked increase in the SP2 fraction and a concomitant decrease in the SP3+SP4 macrophage fraction, corresponding with our observation in *Runx3*^**Δ**^ mice and reports on other mouse strains with IBD development (Biswas et al., 2018; Schridde et al., 2017). In accordance with their normal phenotype, the BM mixed chimera replenished mice exhibited a normal RM waterfall resembling that of the WT BM transferred mice (Figure 4B). Yet, despite the normal colon histology in the mixed chimeric recipient mice, flow cytometry analysis comparison revealed differences in the abundance of WT and Runx3^**Δ**^ LP MNP in the mixed chimeric mice. While the mixed chimera WT/CD45.1 MNP consisted of 75% RM and ~20% CD11b^+^ DC, the mixed chimera Runx3^**Δ**^/CD45.2 MNP displayed a reduction of RM to 60% and a concomitant increased proportion of CD11b^+^ DC to 35% (Figure 4C, S3B). Notably, RM and CD11b^+^ DC subsets distribution in chimeric mice (WT/CD45.1 and *Runx3*^**Δ**^/CD45.2) were similar to the corresponding subsets in either WT or Runx3^**Δ**^ BM replenished mice, respectively. However, the mixed chimeric Runx3^**Δ**^ MNP fraction displayed a decrease in Ly6c^+^ colon monocytes, compared to mice reconstituted only with *Runx3*^**Δ**^ BM (Figure S3B). We then analyzed colonic normalized cell number for each subpopulation. Mice reconstituted with Runx3^**Δ**^ BM had a similar RM cell number as mice reconstituted with WT BM (Figure 4D). Contrarily, Runx3^**Δ**^ BM reconstituted mice had a substantially increased number of CD11b^+^ DC and Ly6c^+^ monocytes (Figure 4E and F). Importantly, the mixed chimeric *Runx3*^**Δ**^/CD45.2 mice displayed a significant reduction in CD11b^+^ DC and Ly6c^+^ monocytes cell number, compared to Runx3^**Δ**^ BM reconstituted mice (Figure 4E and F). These results reveal that Runx3-sufficient MNP suppress the influx and propagation of Runx3^**Δ**^ monocytes and CD11b^+^ DC thereby imposing an anti-inflammatory environment, which prevents the induction of colitis by Runx3^Δ^ MNP. This occurrence is reflected by the re-established equilibrium within the MNP compartment of mixed chimeric mice.

### *Runx3*^Δ^ colon RM transcriptome displays an anti-inflammatory to pro-inflammatory switch and colonic MNP maturation defect

The above presented observations revealed that while colon RM and CD11b^+^ DC are formed in the absence of Runx3, *Runx3*^**Δ**^ mice develop colitis. To gain insight into Runx3-dependent transcriptional program controlling intestinal MNP homeostatic function, we analyzed the transcriptomes of WT, Runx3^**Δ**^ RM and CD11b^+^ DCs. RNA was prepared from sorted cecum RM and CD11b^+^ DC (Figure S4A) of 6-8 weeks old mice. Principal component analysis (PCA) of transcriptome data including all the indicated samples, revealed two distinct cell populations corresponding to CD11b^+^ DC and RM (Figure S4B). Analysis of differential gene expression between Runx3^**Δ**^ and WT RM revealed 70 up-regulated and 128 down-regulated genes in Runx3^**Δ**^ (fold-change ≥ 1.5, p-value ≤ 0.05) (Figure 5A, Table S1). Several of these Runx3^**Δ**^ RM DEGs were validated by RT-qPCR (Figure S4C). In addition, flow cytometry analysis confirmed the up-regulation of two surface proteins encoded by Runx3^**Δ**^ RM DEGs *Pdcd1lg2* and *Clec12a* (Figure S4D, E). Interestingly, gene ontology (GO) enrichment analysis of these DEGs using the term “disease and disorder” yielded “experimental colitis” as the top enriched term in "inflammatory/auto-immune disease" category (Figure 5B). In line with the GO enrichment for the term "colitis", expression of inflammatory genes was affected; 19 out of 21 pro-inflammatory DEGs (85%) were up-regulated and 14 out of 19 (74%) anti-inflammatory DEGs were down-regulated, respectively, in Runx3^**Δ**^ RM (Table 1). These results indicate that loss of Runx3 in MNP switches colonic RM from an anti-inflammatory to a pro-inflammatory state. Of note, GO analysis to detect potential upstream regulators of Runx3^**Δ**^ RM DEGs highlighted IL10RA and IFNG as the most significant regulators, with IL10RA having a negative z-score and IFNG a positive z-score (Figure 5C). Conditional deletion of *Il10ra* in MNP (*Cx3cr1-IL10ra^Δ^*) (Zigmond et al., 2014) and *Il10rb^−/−^* mice (Redhu et al., 2017; Zigmond et al., 2014) develop spontaneous colitis, similar to that observed in *Runx3*^**Δ**^ mice. To examine a possible relationship between Runx3 and Il10 signaling in RM, we cross analyzed Runx3^**Δ**^ DEGs with the transcriptional profile of Il10ra^Δ^ and Il10rb^−/−^ RM. Remarkably, 23 out of the 70 (33%) Runx3^**Δ**^ RM up-regulated genes were also up-regulated in the two other data sets and 40-50% of the Runx3^**Δ**^ RM up-regulated genes overlapped with each of the Il10r^Δ^ RM data sets (Figure 5D). Moreover, the common up-regulated genes in Runx3^**Δ**^, Cx3cr1-Il10ra^Δ^ and Il10rb^*-/-*^ RM included 15 of the 21 (71%) pro-inflammatory genes underscored above (Table 1). While MNP-*Runx3*^**Δ**^ and MNP-*Il10R*^**Δ**^ mice display a similar spontaneous colitis phenotype with a significant overlap of RM up-regulated genes, expression of *Il10ra* and *Il10rb* was unaffected in Runx3^**Δ**^ RM. Together, these observations led us to hypothesize that Runx3 is involved in MNP transcriptional regulation downstream of Il10-induced signaling.

**Figure 5.**
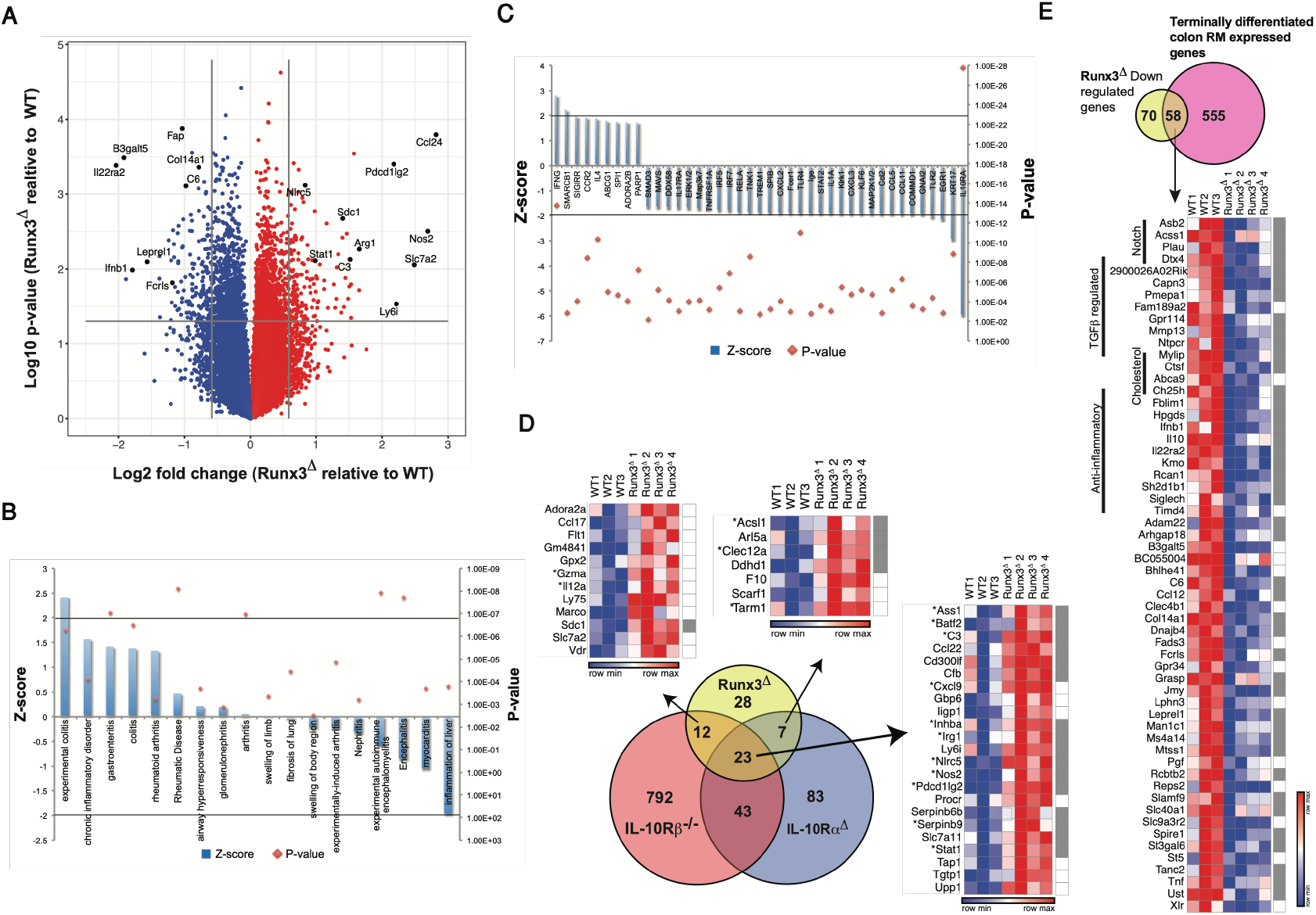
Runx3^**Δ**^ RM transcriptome reveals impaired maturation and up-regulation of pro-inflammatory genes as occurs in Il10r^**Δ**^ RM. **A**, Volcano plot of colon RM DEGs in 6-8 weeks old *Runx3*^**Δ**^ and WT control mice. Numerous up- or down-regulated genes in Runx3^**Δ**^ are indicated. **B**, GO analysis categorized by “disease and disorder”. **C**, GO analysis categorized by “upstream regulators”. **D**, Venn diagram (lower part) representing the overlap between Runx3^**Δ**^, Cx3cr1-Il10ra^**Δ**^ and Il10rb^−/−^ RM up-regulated genes. Cutoff for Runx3^**Δ**^ and the two Il10r-deficient RM DEGs was set to 1.5 and 2-fold change, respectively. Heat maps representing the common genes standardized expression values, asterisks mark the pro-inflammatory genes. Runx3 high-confidence targets are represented by right column grey squares **E**, Venn diagram representing the overlap between down-regulated genes in *Runx3*^**Δ**^ RM and up-regulated genes in terminally differentiated SP4 RM vs. their SP1 monocyte precursors (top). Shared genes standardized expression values are specified in the heat map (bottom). Runx3 high confidence targets are represented by right column grey squares.

**Table 1.**
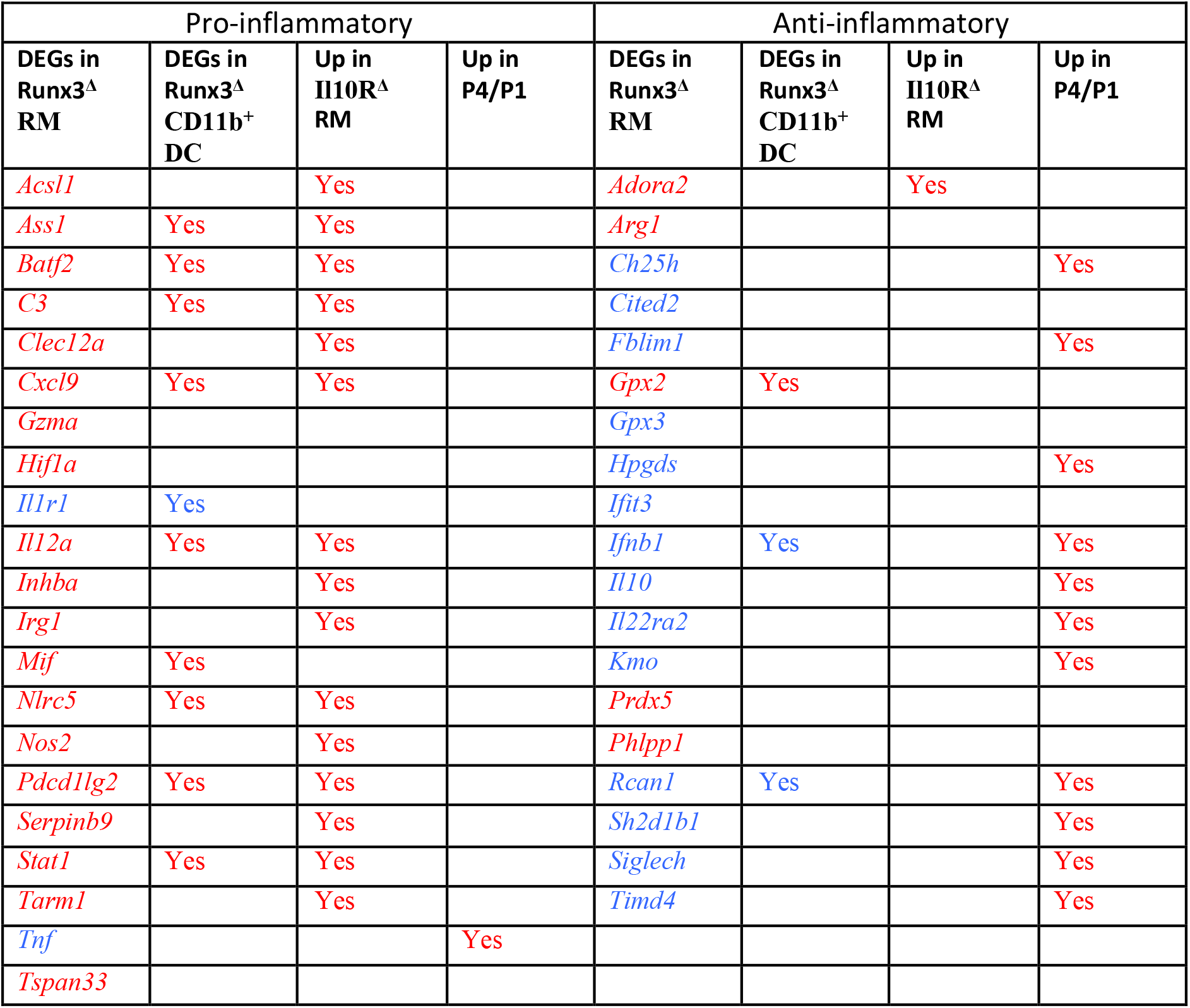
Pro- and anti-inflammatory DEGs in Runx3^Δ^ RM and CD11b^+^ DC and their up-regulation in Il10r^Δ^ and P4 RM. Red and blue marked gene names indicate up-regulated and down-regulated genes, respectively.

Newly arrived colonic monocytes, termed SP1 monocytes, differentiate through stages SP2 and SP3 until ultimately becoming mature SP4 resident macrophages (Schridde et al., 2017). This process is accompanied by expression of 613 genes specifically in SP4 compared to SP1 monocytes (Schridde et al., 2017). As Runx3 is expressed in colonic RM and in its absence the SP4 mature stage is impaired, we hypothesized that Runx3 is involved in the maturation of colon SP4 RM. As mentioned above, 128 down-regulated DEGs were identified by cross analysis of Runx3^**Δ**^/WT RM gene expression profiles (Figure 5A, Table S1). Intersecting these 128 RM Runx3^**Δ**^ down-regulated genes with the 613 SP4-specific genes revealed a marked overlap of 58 genes, comprising 45% of Runx3^**Δ**^ down-regulated genes (Figure 5E). Remarkably, 11 of the 14 (79%) down-regulated anti-inflammatory genes in Runx3^**Δ**^ RM, including *Il10*, were among these 58 common genes (Table 1, Figure 5E). Additionally, among these 58 down-regulated genes several are associated with the Notch pathway and cholesterol uptake. Maturation into colon SP4 RM stage is dependent on up-regulation of various TGFβ signaling genes (Bain et al., 2013; Schridde et al., 2017). As Runx3 has an established role in mediating TGFβ signaling (Fainaru et al., 2004), it is tempting to speculate that the 58 down-regulated genes include Runx3-responsive TGFβ-regulated genes. Cross analysis of the SP4-specific RM gene list with those that were down-regulated in *Tgfbr1*^Δ^*/RAG1*^−/−^ (Schridde et al., 2017), revealed 62 common genes (Table S1). Intersecting these 62 genes with the above mentioned 58 P4-specific genes that were down-regulated in Runx3^**Δ**^ RM, revealed 9 common genes (Figure 5E, Table S1).

To gain insight into potential Runx3 direct target genes, the list of 198 DEGs in Runx3^**Δ**^ RM (Table S1) was cross analyzed with lists of genes containing the hallmarks of transcriptional active regions: ATAC-seq, H3K4me1 and H3K27Ac ChIP-seq peaks in colonic RM (Lavin et al., 2014). Of particular relevance to our findings is the observation that a RUNX motif is highly enriched in enhancer regions specifically in intestinal macrophages as compared to macrophages from all other tissues (Lavin et al., 2014). Interestingly, this analysis revealed that out of these 198 DEGs, 120 (60%) harbored all three peak categories and 28 additional DEGs harbored both H3K4me1 and H3K27Ac peaks (Figure S5A), raising the number of peak-bearing DEGs to 148 (75%). Accordingly, only 25 DEGs (13%) did not contain any of these peaks (Figure S5A). Moreover, 126 of these 148 peak-harboring genes contained at least one region in which at least two of the three above mentioned peak categories overlapped (in most cases H3K4me1 and H3K27ac). Manual analysis of the 126 DEGs containing overlapping peaks, including the 40 pro- and anti-inflammatory DEGs (Table 1), revealed that 117 of them (95%) harbored a RUNX motif in the overlapping peak (Table S1 and see *Ass1* and *Il10* as examples in Figure S5B). We defined these Runx3^**Δ**^ RM DEGs as high-confidence Runx3 targets. Interestingly, the human homologs of 8 of these RM Runx3 high-confidence target genes, *CFB, IFIH1, IL10, NOS2, PLAU, PRDX5 and TNF*, harbor SNPs that are susceptibility loci for IBD, Crohn’s disease (CD) and/or ulcerative colitis (UC) (Table S1). Finally, we also noted that six out of the nine (Figure 5E) Runx3 putative targets regulated by TGF-β in RM SP4 population harbor a RUNX-SMAD module in their Runx3 bound regions (four of these are shown in Figure S5C). Overall, the results suggest that Runx3 is involved in positively regulating TGFβ-dependent RM maturation and Il10-driven anti-inflammatory response while suppressing a pro-inflammatory program.

### Runx3 positively regulates colonic CD11b^+^ DC differentiation genes

The colonic CD11b^+^ cDC2 consist of double positive CD103^+^CD11b^+^ DC and their precursors CD103^+^CD11b^+^ DC. Both DC subsets express Runx3 (Figure 2E). The increased prevalence of colonic LP CD103^+^CD11b^+^ and the decrease in CD103^+^CD11b^+^ DC in *Runx3*^*Δ*^ compared to WT mice (Figure 3C) raised the possibility that, as in RM, Runx3 affects CD103^+^CD11b^+^ DC differentiation. Because it was hard to isolate sufficient numbers of colonic CD103^+^CD11b^+^ DC from *Runx3*^*Δ*^ mice, we compared the transcriptome of WT and Runx3^*Δ*^ colonic CD11b^+^ DC, including both CD103^+^CD11b^+^ and CD103^+^CD11b^+^ DC. The transcriptome profile revealed 84 up-regulated and 152 down-regulated genes (fold-change ≥ 1.5, p-value ≤ 0.05) in Runx3^*Δ*^ vs. WT CD11b^+^ DC (Figure 6A, Table S1). Similar to RM, GO analysis of DEGs in CD11b^+^ DC underscored IL10RA and IFNG as the top significant upstream regulators (Figure 6B). Importantly, we found a subset of 31 Runx3^**Δ**^ DEGs common to both RM and CD11b^+^ DC (Figure 6C), suggesting that some Runx3-regulated functions are shared between these two MNP populations. Remarkably, 13 of these 31 Runx3^**Δ**^ RM and CD11b^+^ DC common DEGs (42%) are inflammation-regulating genes, including 10 up-regulated pro-inflammatory and 3 down-regulated anti-inflammatory genes (Table 1). Moreover, most of the pro-inflammatory genes up-regulated in Runx3^**Δ**^ CD11b^+^ DC were shared by Il10ra and/or Il10rb-deficient RM (Table 1). These results suggest that Runx3^**Δ**^ CD11b^+^ DC display pro-inflammatory properties, similar to monocyte-derived CD11b^+^ DC following mild DSS-induced colitis (Varol et al., 2009). In addition, Runx3^**Δ**^ CD11b^+^ DC showed down-regulation of *Ifnb1* and IFN-β-regulated genes *Ifih1* and *Mx2* (Figure 6C, Table S1). Furthermore, enrichment of genes associated with β-catenin and TGF-β signaling was noted among Runx3^**Δ**^ CD11b^+^ DC down-regulated genes (Figure 6D).

**Figure 6.**
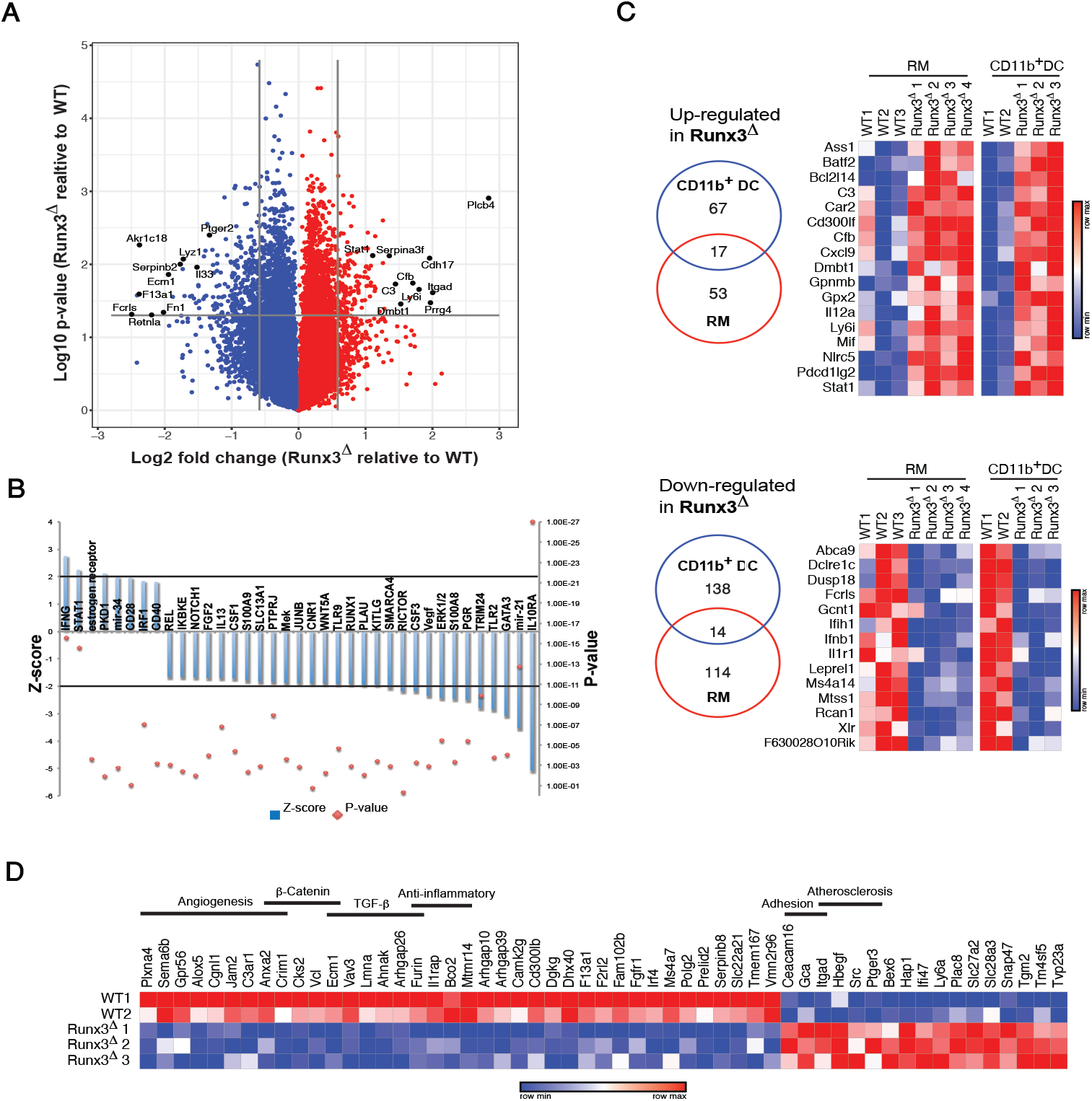
A fraction of DEGs between Runx3^*Δ*^ and WT CD11b^+^ DC are shared with Runx3^*Δ*^ RM. **A**, Volcano plot of colon CD11b^+^ DC DEGs in 6-8 weeks old *Runx3*^**Δ**^ and WT control mice. Some up- or down-regulated genes in Runx3^**Δ**^ CD11b^+^ DC are indicated. **B**, GO analysis categorized by up-stream regulators. Cut-off and z-score values were set to 2. C, Venn diagrams and heat maps showing common DEGs in colonic Runx3^**Δ**^ CD11b^+^ DC and RM. **D**, Heat map comprising standardized expression values of 65 high-confidence Runx3 target genes in CD11b^+^ DC.

Runx3 ChIP-seq analysis in the D1 DC cell line and splenic CD4^+^ DC (Dicken et al., 2013) revealed 13,014 and 15,121 Runx3-bound regions (peaks), of which 6836 peaks overlapped, corresponding to 6422 genes. Interestingly, GREAT analysis under “PANTHER Pathway” revealed that the genes corresponding to the overlapping Runx3-bound peaks in D1 cells and splenic CD4^+^ DC, as well the genes corresponding to the peaks in colonic RM (Lavin et al., 2014), are highly enriched for the term “Inflammation mediated by chemokine and cytokine signaling pathway” (Table S1). To determine the putative CD11b^+^ DC direct Runx3 target genes, we first cross-analyzed the list of DEGs in colonic Runx3^**Δ**^ CD11b^+^ DC with the list of common Runx3-bound genes in D1 and splenic CD4^+^CD11b^+^ DC, revealing that 90 of the 236 DEGs in Runx3^**Δ**^ CD11b^+^ DC (38%) harbored Runx3-bound regions (Figure S6A). Moreover, 65 of these 90 Runx3-bound DEGs (71%) contained at least one region with a RUNX motif, suggesting that they are high-confidence Runx3-target genes (Figure 6D, Table S1). Moreover, 10 of the 31 common DEGs in Runx3^**Δ**^ CD11b^+^ DC and RM (32%) are also common high-confidence Runx3 target genes (Figure 6C and S6B). Interestingly, 3 of these 10 common Runx3 target genes in colonic RM and CD11b^+^ DC (*Ifnb1*, *Pdcd1lg2* and *Stat1*) are pro- or anti-inflammatory genes (Figure S6C), suggesting that they might contribute substantially to *Runx3*^**Δ**^ mice colitis phenotype. Of note, the human homologs of 4 of the CD11b^+^ DC Runx3 targets, *CD300LF, IFIH1*, IRF4 and *SLC22A5* (mouse *Slc22a21*) harbor SNPs associated with IBD, CD, UC and/or celiac (Table S1) and the 2 former genes are common Runx3 targets in RM and CD11b^+^ DC. Overall, the results indicate that Runx3 is important for the maturation of CD11b^+^ DC into anti-inflammatory and tolerogenic DC and loss of Runx3 in CD11b^+^ DC affects these anti-inflammatory and tolerogenic properties in a remarkably similar way to that found in Runx3^*Δ*^ RM and Il10r deficient RM.

### Runx3 loss in MNP induces tolerogenic to inflammatory CD4 T-cell transition

The MNP switch from anti-to pro-inflammatory response in the absence of Runx3 implied impaired activation of T cells. Altered expression of Tregs inducing genes further supported this possibility. Two down-regulated genes in cDC2, *Il33* and *Il6*, encode important cytokines for intestinal Tregs generation (Schiering et al., 2014), (Table S1). Furthermore, it was recently shown that an additional cDC2 down regulated gene, IFN-β is important in intestinal control of Tregs differentiation (Nakahashi-Oda et al., 2016). To verify if Tregs are affected, we compared colonic Foxp3^+^ Tregs of WT and *Runx3*^**Δ**^ mice. The prevalence of Tregs was determined by gating on CD45^+^CD4^+^CD45RB^lo^ cells, followed by gating on CD25^+^Foxp3^+^ cells. Strikingly, Runx3^**Δ**^ colon LP showed a substantial reduction in the frequency of Foxp3^+^ Tregs (Figure 7A).

**Figure 7.**
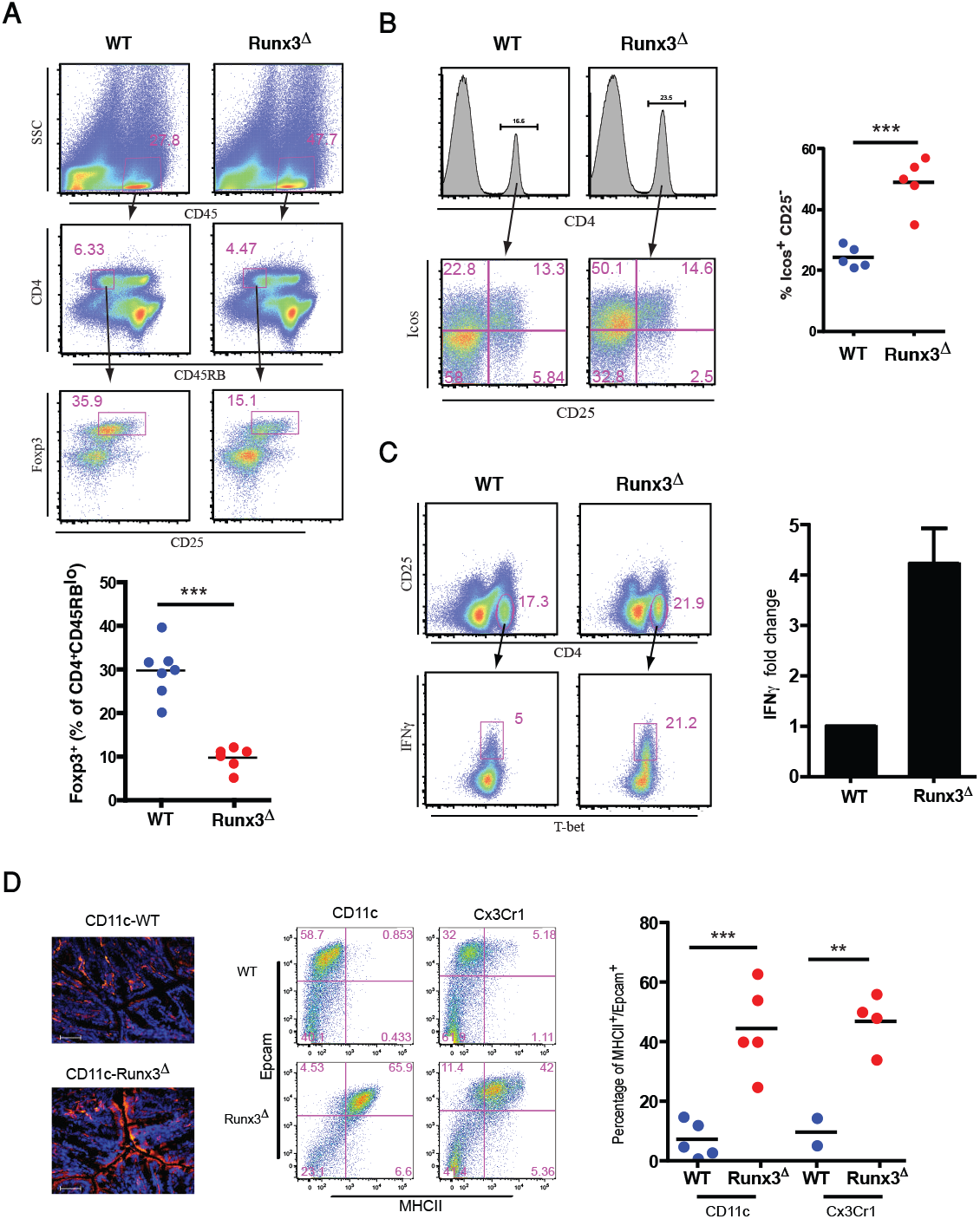
Runx3-deficient MNP cause a tolerogenic to inflammatory switch in colonic CD4^+^ T cells. **A**, Analysis of Tregs in *Runx3*^**Δ**^ colonic LP. Representative flow cytometry analysis (top) and graphical summary (bottom) of CD25^+^Foxp3^+^ Tregs in colonic LP of WT *MNP-Runx3*^**Δ**^ mice. ***P<0.001. **B**, Representative flow cytometry (left) and graphical summary (right) of Icos-expressing WT and Runx3^**Δ**^ colonic LP CD4^+^CD25^−^ T cells. **C**, Intracellular staining of IFN-g in WT and Runx3^**Δ**^ colonic LP CD4^+^CD25^−^ T cells. **D**, MHCII IF staining in WT and *Runx3*^**Δ**^ colon epithelium. Frozen sections from WT and *Runx3*^**Δ**^ colons stained with anti-MHCII. Magnification x40; scale bars 50 pm (left). Flow cytometry analysis (middle) and graphical summary (right) of colonic epithelial cells from WT, *Runx3*^**Δ**^ and *Cx3cr1-Runx3*^**Δ**^ mice stained with anti-EpCAM and anti-MHCII Phycoerythrin (PE). * **P<0.01, ***P<0.001.

The inducible T-cell co-stimulator (ICOS) plays role in modulating different adaptive immune responses. Hence, we examined whether ICOS expression is affected in CD4^+^ lymphocytes of *Runx3*^**Δ**^ mice. Interestingly, we found that in *Runx3*^**Δ**^ mice the abundance of ICOS expressing cells in CD4^+^ lymphocytes is significantly increased compared to WT (Figure 7B) suggesting an increase of activated CD4^+^ T cells in the *Runx3*^**Δ**^ mice. GO analysis results suggested that Runx3^**Δ**^ MNP might also induce T-cell activation manifested by IFN-γ production. To examine the impact of Runx3 deletion in MNP on lymphocyte activation we analyzed IFN-γ expression in LP CD4^+^ T cells treated with TPA and ionomycin. The results indicate increased abundance of IFN-γ expressing activated LP CD4^+^ T-cells in *Runx3*^**Δ**^ mice (Figure 7C). Mucosal immunity depends partly on cross talk between the immune system and the epithelium. For instance, IFN-γ produced by CD4^+^ T cells plays a role in ameliorating colitis development by induction of MHCII expression in epithelium (Mayer et al., 1991; Thelemann et al., 2014). Induction of MHCII in mucosal epithelial cells was also reported in IBD patients (Buning et al., 2006). Consequently, we addressed whether colonic epithelium MHCII is affected in *Runx3*^**Δ**^ mice. Analysis revealed a profound adluminal MHCII expression in Runx3^**Δ**^ LP epithelial cells, whereas MHCII expression in the WT LP was confined to leukocytes (Figure 7D left). Furthermore, we detected MHCII expression in colonic epithelium in both *Runx3*^**Δ**^ and *Cx3cr1-Runx3^Δ^* mice, as evidenced by co-expression of MHCII and epithelial marker EpCAM (Figure 7D middle and right). Collectively, the results imply that Runx3 expression in colonic LP MNP plays an important role in promoting intestinal immunological tolerance, reflected by induction of LP Foxp3^+^ Tregs and reduction in IFN-γ producing CD4^+^ T-cells. The diversion from regulatory into pro-inflammatory CD4^+^ T cells in Runx3^**Δ**^ colon reinforces the impact of the transcriptional transition of Runx3^**Δ**^ MNP into an immature pro-inflammatory state.

## DISCUSSION

The large surface area of the intestinal mucosa exposed to the external environment poses a great challenge to the mucosal immune system to balance between defense against pathogens and tolerance to commensal bacteria and food antigens. Breaking this balance leads to IBD (Gross et al., 2015). More than 200 human IBD susceptibility loci were identified, mostly in genomic regions in the vicinity of genes expressed in various immune cells, including on chromosome 1p36 in the *RUNX3* locus (Cho, 2001; Cho et al., 1998; Lotem et al., 2017; Peters et al., 2017). Moreover, many IBD susceptibility loci were associated with genes that are RUNX3 targets in various immune cells (Lotem et al., 2017). Previously we reported that Runx3^−/−^ mice develop early onset spontaneous colitis (Brenner et al., 2004). The fact that Runx3 is not expressed in normal colonic epithelium (Brenner et al., 2004; Levanon et al., 2011), suggested that loss of a leukocytic cell-autonomous Runx3 function was the driving force of colitis development in these mice (Brenner et al., 2004). Here we provide direct evidence that transfer of Runx3^−/−^, but not WT, FL or BM cells into lethally irradiated mice induced colitis in the recipient mice. Moreover, conditional deletion of *Runx3* in MNP, but not in lymphocytes, recapitulated the spontaneous development of colitis observed in *Runx3^−/−^* mice. These results indicate that Runx3 expression in MNP is important for their known role in maintaining GIT homeostasis (Joeris et al., 2017).

Runx3 loss in MNP results in a decreased abundance of colonic SP3-SP4 mature anti-inflammatory RM and CD103^+^CD11b^+^ cDC2, which occurs prior to the onset of significant colitis symptoms, suggesting that these changes are indeed the cause of colitis. This premise is supported by the finding that Runx3^**Δ**^ BM transfer to lethally irradiated mice induced the same changes in MNP populations as well as colitis, whereas mice transplanted with an equal number of WT and Runx3^**Δ**^ BM cells showed a normal MNP balance and remained healthy. These data are consistent with the conclusion that presence of WT BM cells overcomes the colitogenic effect of the transplanted Runx3^**Δ**^ BM cells.

Analysis of Runx3^**Δ**^ and WT colonic RM transcriptomes, revealed a prominent up-and down-regulation of pro- and anti-inflammatory genes, respectively, in Runx3^**Δ**^ RM. These results strongly indicate that loss of Runx3 in RM induces an anti-to pro-inflammatory switch in their biological properties. Comparison of RM DEGs in Runx3^**Δ**^ with those in Il10ra^**Δ**^ and Il10rb^−/−^, that also induce colitis (Redhu et al., 2017; Zigmond et al., 2014), revealed gain of pro-inflammatory hallmarks in all three models of spontaneous colitis. Loss of Runx3 in RM did not affect expression of *IL10ra* and *IL10rb* in our data and neither was expression of *Runx3* significantly affected by loss of Il10rb (Redhu et al., 2017). Therefore, the similar spontaneous colitis phenotype in these three strains cannot be explained by cross-regulation between Runx3 and signals emanating from the Il10 receptors. However, it is possible that activation of Stat3 TF, downstream of Il10 receptor signaling (Biswas et al., 2018) collaborates with Runx3 in the nucleus, which can explain the partially shared effect on gene expression when either of these TFs is deleted. This possibility is supported by the known ability of Stat and Runx proteins to physically interact (Ogawa et al., 2008) and the spontaneous colitis induced in mice harboring Stat3-deficient MNP (Kobayashi et al., 2003; Melillo et al., 2010; Takeda et al., 1999). Of note, the fact that the common DEGs in Runx3- and Ill0 receptor-deficient RM is mostly confined to up-regulated genes, suggests that the collaboration between Runx3 and Stat3 is employed mainly to suppress their targets, notably pro-inflammatory genes. It should also be stressed that while Runx3 and Il10R deficiencies in RM lead to colitis, each model bears unique features. For example, down regulation of anti-inflammatory genes occurs in Runx3^**Δ**^ RM, but is neglected in the two Il10r models. Furthermore, it was recently reported that Il10ra^**Δ**^ RM show increased expression of Il23, which induces Il22 production in T cells leading to hypertrophy of colonic epithelium (Bernshtein et al., 2019). In contrast, there was no change in *Il23* expression in Runx3^**Δ**^ RM, but expression of *Il22ra2*, encoding a very potent antagonist of Il22 receptor signaling, was down-regulated. Because the balance of Il22 and its antagonist Il22ra2 (IL22RB) is important to maintain intestinal homeostasis (Zenewicz, 2018), it is conceivable that reduced *Il22ra2* expression in Runx3^**Δ**^ RM, increases response to Il22 itself and could thus elicit an inflammatory response in the epithelium.

The differentiation of intestinal SP1 monocytes to fully mature antiinflammatory SP4 RM is a TGFβ-dependent process (Schridde et al., 2017) and we have now shown that SP4 RM expressed a higher level of Runx3 compared to SP1 monocytes. Loss of Runx3 in MNP results in defective RM differentiation associated with reduced expression of anti-inflammatory genes. The finding of reduced expression of TGFβ and Notch-regulated genes is in-line with the defect of Runx3^**Δ**^ RM differentiation. Thus, Runx3 function in normal RM to repress pro-inflammatory and induce anti-inflammatory genes, is consistent with its ability to protect against colitis. Finally, ~60% of all DEGs in Runx3^**Δ**^ RM, including most of the inflammatory genes, are in fact high-confidence Runx3-target genes as judged by their harboring overlapping ATAC and enhancer chromatin marks peaks containing a RUNX binding motif. Moreover, as with human *RUNX3* itself (Lotem et al., 2017), the human homologs of 8 of these high-confidence Runx3 target genes in RM contain known susceptibility loci for IBD, CD, UC or celiac GIT diseases.

Runx3 is a key player in cell-lineage fate decisions including in DC and do so to some extent by mediating response to TGFβ. For example, Runx3 regulates TGFβ-mediated lung DC cell functions, facilitates specification of murine splenic CD11b^+^Esam^hi^ DC and is mandatory to the TGFβ-dependent development of skin Langerhans cells (Dicken et al., 2013; Fainaru et al., 2004). TGFβ-dependence of differentiation of intestinal CD103^+^CD11b^+^ DC was also demonstrated (Bain et al., 2017; Schridde et al., 2017). We found that Runx3 is expressed at low levels in ~ 20% of intestinal cDC1 whereas it is highly expressed in the majority of cDC2 cells. Given that Runx3^**Δ**^ LP shows reduced abundance of these mature CD103^+^CD11b^+^ cDC2, it is conceivable that Runx3 participates in TGFβ signaling in cDC2. Mice lacking colonic cDC1 can still establish tolerance, presumably by their CD11b^+^ cDC2 (Veenbergen et al., 2016), so the fact that colonic CD11b^+^ cDC2 are affected in *Runx3*^**Δ**^ mice may imply that these cells contribute to colitis development. Analysis of cDC2 transcriptomes of Runx3^**Δ**^ vs. WT revealed that merely 31 of the 236 DEGs in Runx3^**Δ**^ cDC2 were common with those of Runx3^**Δ**^ RM. Remarkably, ~40% of these 31 common DEGs were inflammation-regulating genes, including 10 commonly up-regulated pro-inflammatory genes and 3 commonly down-regulated anti-inflammatory genes. Most of these common pro-inflammatory genes up-regulated in Runx3^**Δ**^ RM and cDC2 are also up-regulated in Il10ra and/or Il10rb-deficient RM (Table 1). These results imply that like their Runx3^**Δ**^ RM counterparts, LP Runx3^**Δ**^ cDC2 contribute to the colitis phenotype by acquiring a pro-inflammatory state. Cross-analysis of Runx3^**Δ**^ cDC2 DEGs with genes that harbored overlapping Runx3-bound peaks in ChIP-seq assays revealed that 65 of them contained a RUNX motif marking them as high-confidence Runx3 target genes. Ten of these cDC2 high-confidence Runx3 targets were common to those in RM and the human homologs of 4 of them *CD300LF, IFIH1, IRF4* and *SLC22A5* (mouse *Slc22a21*) harbor SNPs associated with IBD, CD, UC and/or celiac. Of particular interest is the known importance of Irf4 for the survival of intestinal CD103^+^CD11b^+^ DC (Persson et al., 2013), which can explain the reduced abundance of these cells in Runx3^**Δ**^ colon.

Intestinal DCs participate in immune tolerance and barrier protection by driving differentiation of regulatory T cells and Th17 cells, respectively (Zhou and Sonnenberg, 2018). Interestingly, some of DEGs in Runx3^**Δ**^ cDC2 suggest an impairment of the β-catenin signaling pathway, which is important in the induction of tolerogenic DC (Bain et al., 2013; Schridde et al., 2017). Another critical element in the tolerance breach in Runx3^**Δ**^ mice is the substantial reduction in Foxp3^+^ regulatory T cells, a crucial component in induction of GIT mucosal tolerance. This reduction can be attributed to a defective ability of Runx3^**Δ**^ cDC2 to generate regulatory T cells since essential Treg-inducing factor genes such as *Aldh1a2*, *Il33* and *Ifnβ* (Bakdash et al., 2015; Nakahashi-Oda et al., 2016; Schiering et al., 2014) were down regulated in Runx3^**Δ**^ cDC2. The decrease in *Ifnβ* expression also in Runx3^**Δ**^ RM suggests that loss of Runx3 expression in RM participates in the failure of Runx3^**Δ**^ mice to generate and/or maintain GIT Tregs. In addition, the induced surface expression of Pd-l2 in Runx3^**Δ**^ MNP exemplifies the pro-inflammatory state acquired by Runx3^**Δ**^ RM, which can explain the increased production of IFNγ by Runx3^**Δ**^ CD4^+^ T-cells.

Besides the similarity in inflammatory DEGs between Runx3^**Δ**^ RM and cDC2, other DEGs unique to each MNP subset may also contribute to the colitis phenotype. RM are normally maintained in an inflammatory anergy state by acquisition of a non-inflammatory gene expression profile (Bain et al., 2013; Rivollier et al., 2012; Weber et al., 2011), yet they retain their bactericidal capacity (Niess et al., 2005). Interestingly, the anti-bacterial autophagy gene *Clec12a* reported to be associated with an increased risk for CD (Begun et al., 2015) and *Slc7a11*, a potential blocking target for treatment of IBD patients (Bridges et al., 2012), showed up-regulation specifically in Runx3^**Δ**^ RM but not in Runx3^**Δ**^ cDC2.

While the increased and decreased expression of pro-inflammatory and anti-inflammatory genes, respectively, in Runx3^**Δ**^ MNP can contribute to the development of colitis, it may not be sufficient, since in the context of the BM chimera replenishment assays the mixed WT/Runx3^**Δ**^ MNP chimeric mice were protected from colitis. This finding implies that WT Runx3-suficient MNP confer an immune-suppressive GIT condition, by maintaining a proper balance of pro- and anti-inflammatory gene expression in MNP themselves together with an indirect effect that prevents loss of Tregs. To summarize, all our results point towards one major conclusion: MNP Runx3 maintains colon homeostasis by directing proper colon MNP specification into mature anti-inflammatory MNP and concomitantly repressing expression of a harmful pro-inflammatory program, similar to that which occurs in Il10 receptor-deficient MNP. Another layer of MNP Runx3 contribution to intestinal homeostasis is by its impact on maintaining colonic Tregs. These results imply that human MNP RUNX3 plays an important role in preventing development of inflammatory GIT diseases in humans, including IBD, CD, UC and celiac, a premise that is strongly supported by the presence of susceptibility loci for these diseases in the RUNX3 gene itself and in 10 other genes that are high-confidence RUNX3 targets in RM and/or cDC2.

## ACKNOWLEDGMENTS

We thank Steffen Jung and Ehud Zigmond from the Weizmann Institute Department on Immunology for helpful discussions and for providing the D1 cell line, the Cx3cr1-Cre, Cx3cr1-GFP and CD45.1 mice. Ofira Higfa, Rafael Saka and Pavel Bell for animal husbandry, Sima Peretz for assistance in intravenous injections and Calanit Raanan for tissue sections preparation and Shirely Horn Saban for help in gene expression data acquisition.

## AUTHOR CONTRIBUTIONS

SH, VN, OB, DG, and JD performed experiments. SH, DeL, JL, DiL, OB and YG designed experiments and analyzed data. SH, YL, DL and YG wrote the paper.

The authors declare that they have no conflict of interest

