## Supplemental Information for "Runx3 prevents spontaneous colitis by directing differentiation of anti-inflammatory mononuclear phagocytes"

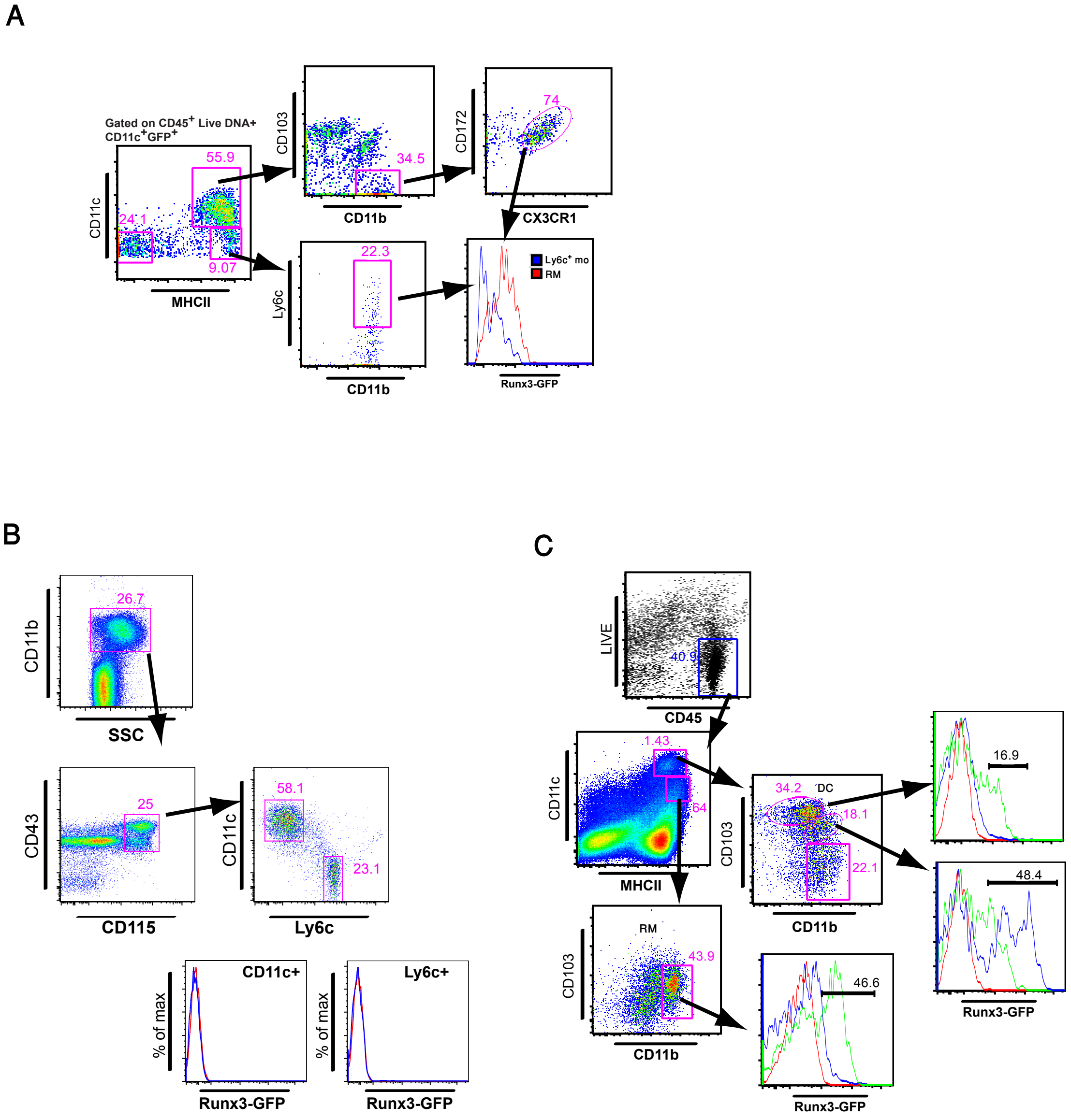

**Figure S1**. **A**, Flow cytometry analysis of Runx3-GFP expression in LP RM and monocytes. **B,** Flow cytometry analysis of Runx3-GFP expression in the two circulating blood monocyte subsets. **C,** Flow cytometry analysis of Runx3-P1$^{AFP/+}$ (blue) or Runx3-P2$^{GFP/+}$ (green) expression in LP RM and CD103$^{+}$CD11b$^{+}$ DC relative to WT control (red). Representative experiment is shown.

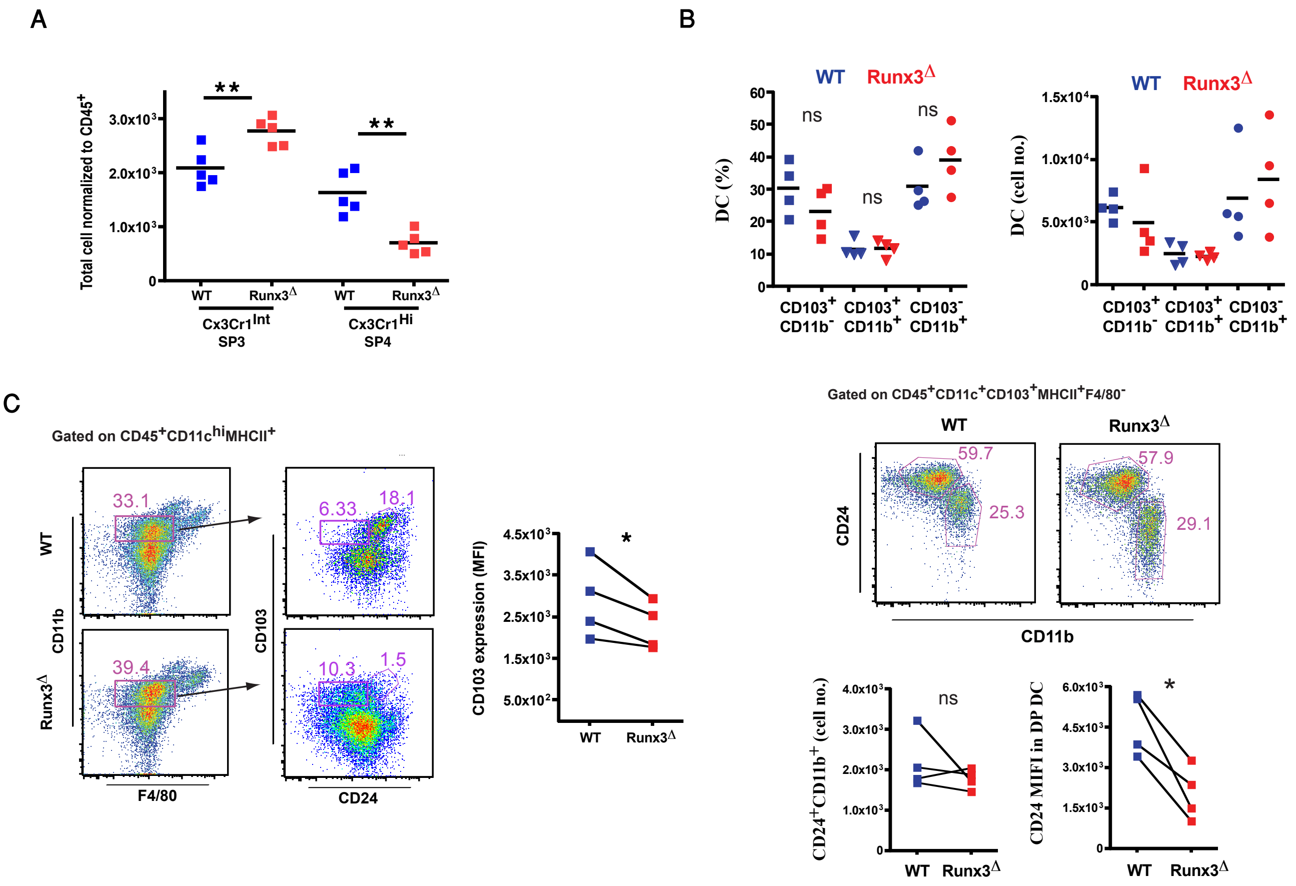

**Figure S2**. **A,** Graphical summary of total number of Cx3cr1$^{Int}$ and Cx3cr1$^{Hi}$ CD11b$^{+}$ RM in *Runx3*$^{\Delta}$-*Cx3cr1*$^{GFP/+}$ and WT-*Cx3cr1*$^{GFP/+}$, **P<0.01. **B,** Graphical summary comparison of DC subsets prevalence and cell subsets number between *Runx3*$^{\Delta}$ and WT at 5-6 weeks (top). Representative flow cytometry comparing CD24a expression on the CD103$^{+}$CD11b$^{+}$ DC. Note the reduced CD24a expression in *Runx3*$^{\Delta}$ (middle). Graphical summary comparing CD103$^{+}$CD11b$^{+}$CD24a$^{+}$ cell number and CD24 expression level between *Runx3*$^{\Delta}$ and WT in 6 weeks old mice. Note the reduced CD24a expression in *Runx3*$^{\Delta}$ (bottom). **C,** Representative flow cytometry comparing CD103 expression on the CD11b$^{+}$ DC. *Runx3*$^{\Delta}$ and WT mice analyzed at 6 weeks of age. Note the reduced CD103 expression in *Runx3*$^{\Delta}$ (middle). Graphical summary comparing CD103 expression level in CD11b$^{+}$ DC between Runx3$^{\Delta}$ and WT (right).

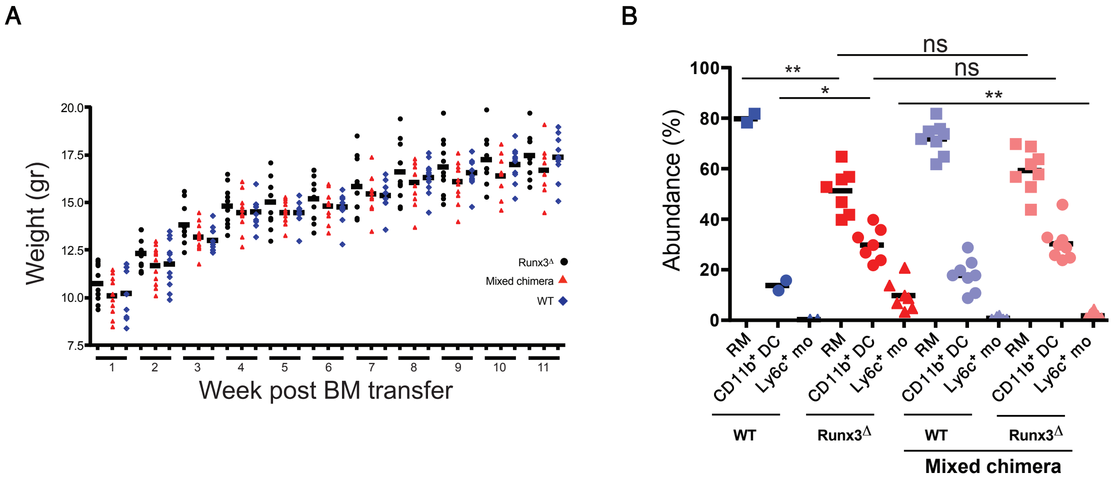

**Figure S3**. **A,** Weight follow-up of Runx3$^{\Delta}$, mixed chimera and WT BM recipient mice. **B,** Abundance of RM, CD11b$^{+}$ DC and Ly6c$^{+}$ monocytes in Runx3$^{\Delta}$, mixed chimera and WT BM recipient mice. * $p<0.05$, ** $p<0.01$.

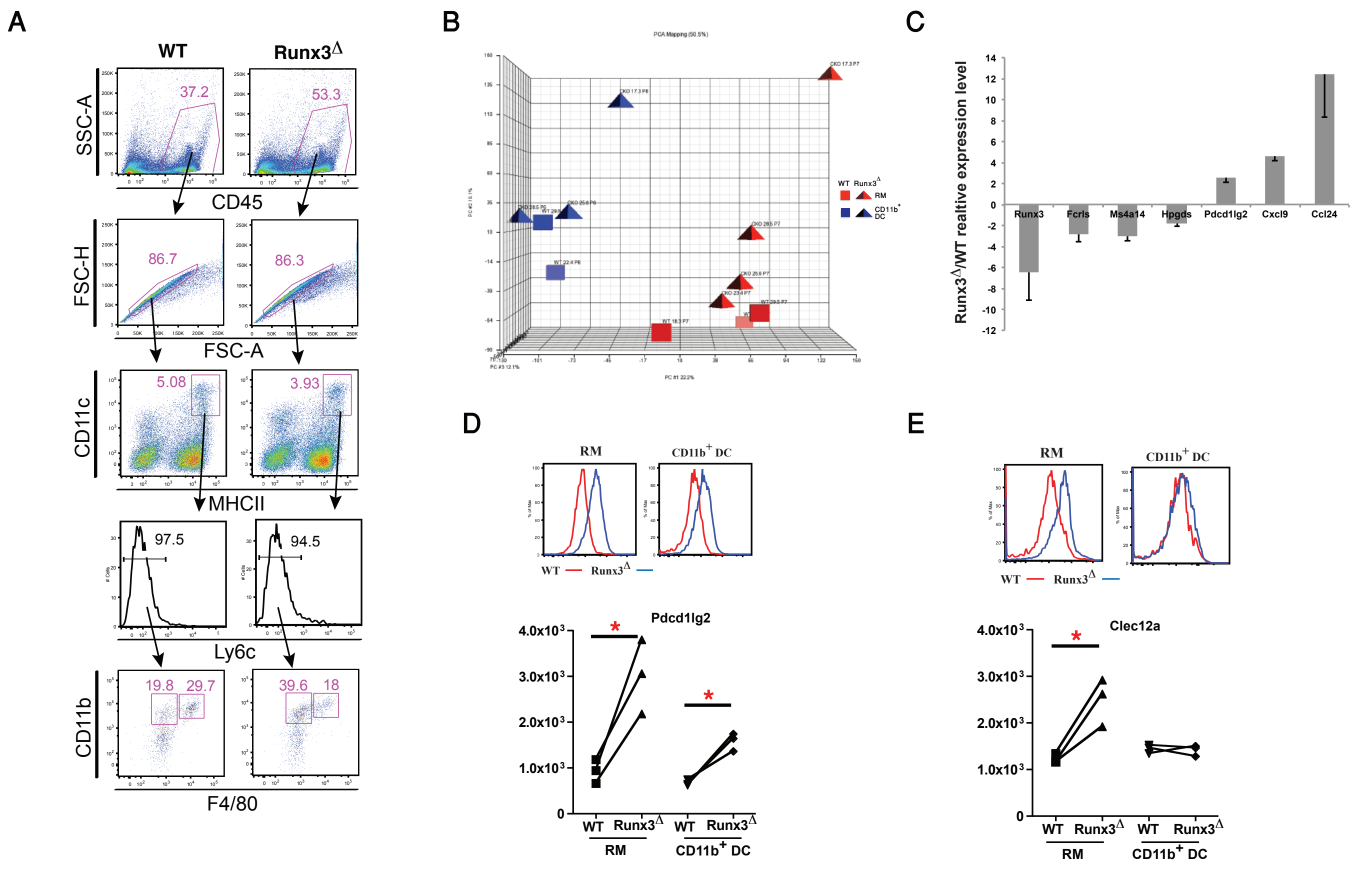

**Figure S4**. Transcriptome analysis of WT vs Runx3$^{\Delta}$ RM and CD11b$^{+}$ DC. **A,** Sorting strategy of colonic LP RM (CD45$^{+}$CD11c$^{+}$MHCII$^{+}$CD11b$^{+}$F4/80$^{+}$Ly6c$^{-}$) and CD11b$^{+}$ DC (CD45$^{+}$CD11c$^{+}$MHCII$^{+}$CD11b$^{+}$F4/80$^{-}$Ly6c$^{-}$) from 6-8 weeks old *Runx3*$^{\Delta}$ and WT littermate mice. Cells sorted from 3-4 mice were pooled in each sample. **B,** PCA analysis of all microarray samples, showing clear separation of RM and CD11b$^{+}$ DC. **C,** qPCR validation of DEGs between Runx3$^{\Delta}$ and WT RM in the microarray analysis. **D,** Comparison of Pdcd1lg2 expression between Runx3$^{\Delta}$ and WT RM and CD11b$^{+}$ DC. Flow cytometry (top) and graphical summary (bottom) of 3 biological repeats are shown. *P<0.05. **E,** Comparison of Clec12a expression between Runx3$^{\Delta}$ and WT RM and CD11b$^{+}$DC. Flow cytometry (upper) and graphical summary (lower) of 3 biological repeats are shown. *P<0.05.

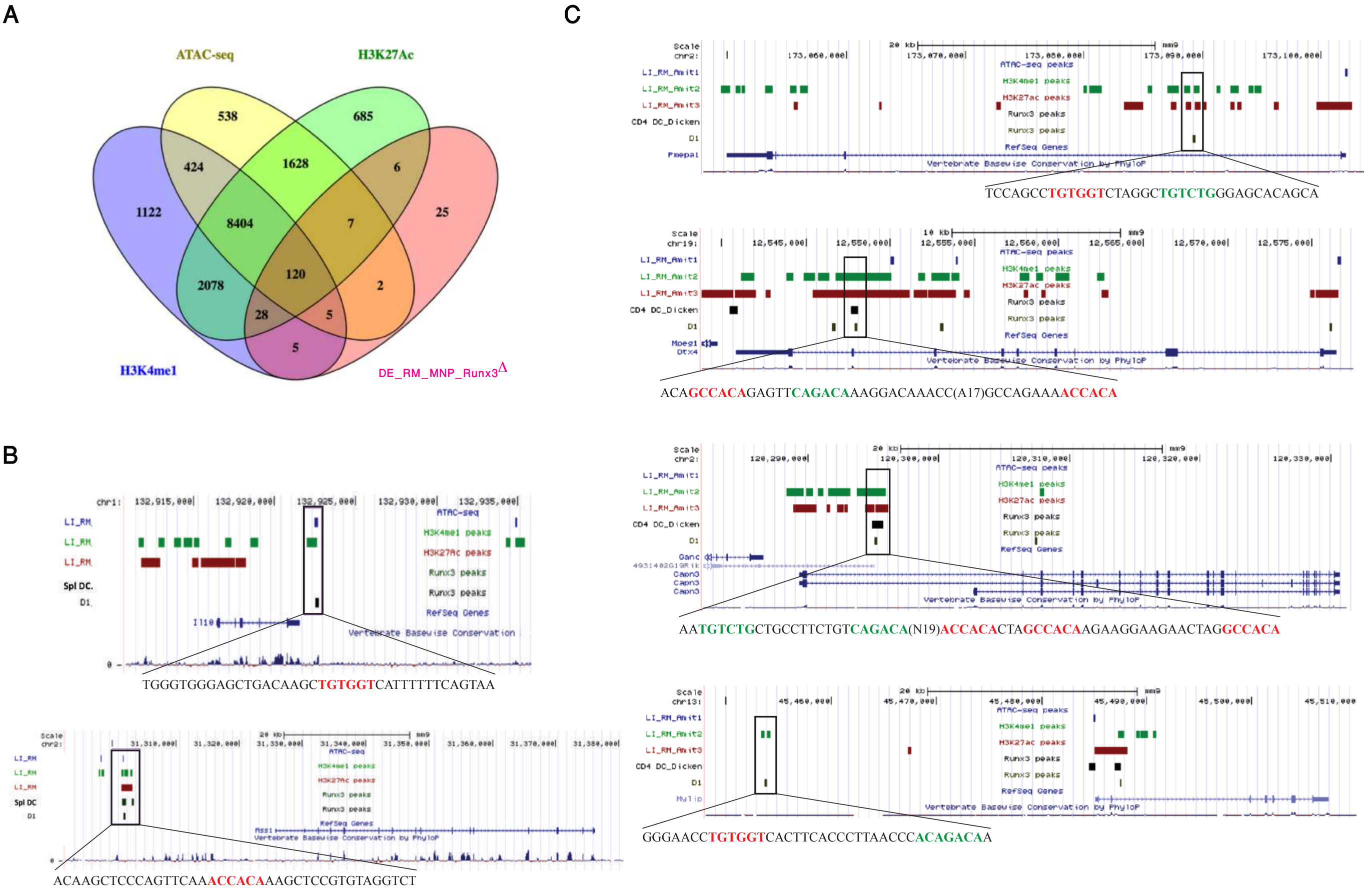

**Figure S5**. **A,** Intersection of DEGs in colonic LP $Runx3^{\Delta}$ RM with genes bearing ChIP-seq peaks in WT colonic RM, splenic CD4$^{+}$ DC and D1 cells. **B,** UCSC genome browser display of two high-confidence DEGs, *Il10* and *Ass1* (top and bottom, respectively), with peaks containing a RUNX motif in the boxed region. **C,** UCSC genome browser display of four TGF-β regulated high-confidence Runx3 target genes in RM. Peaks containing a RUNX and SMAD motifs are marked by boxed region. The DNA sequences demonstrate the RUNX (red)-SMAD (green) module.

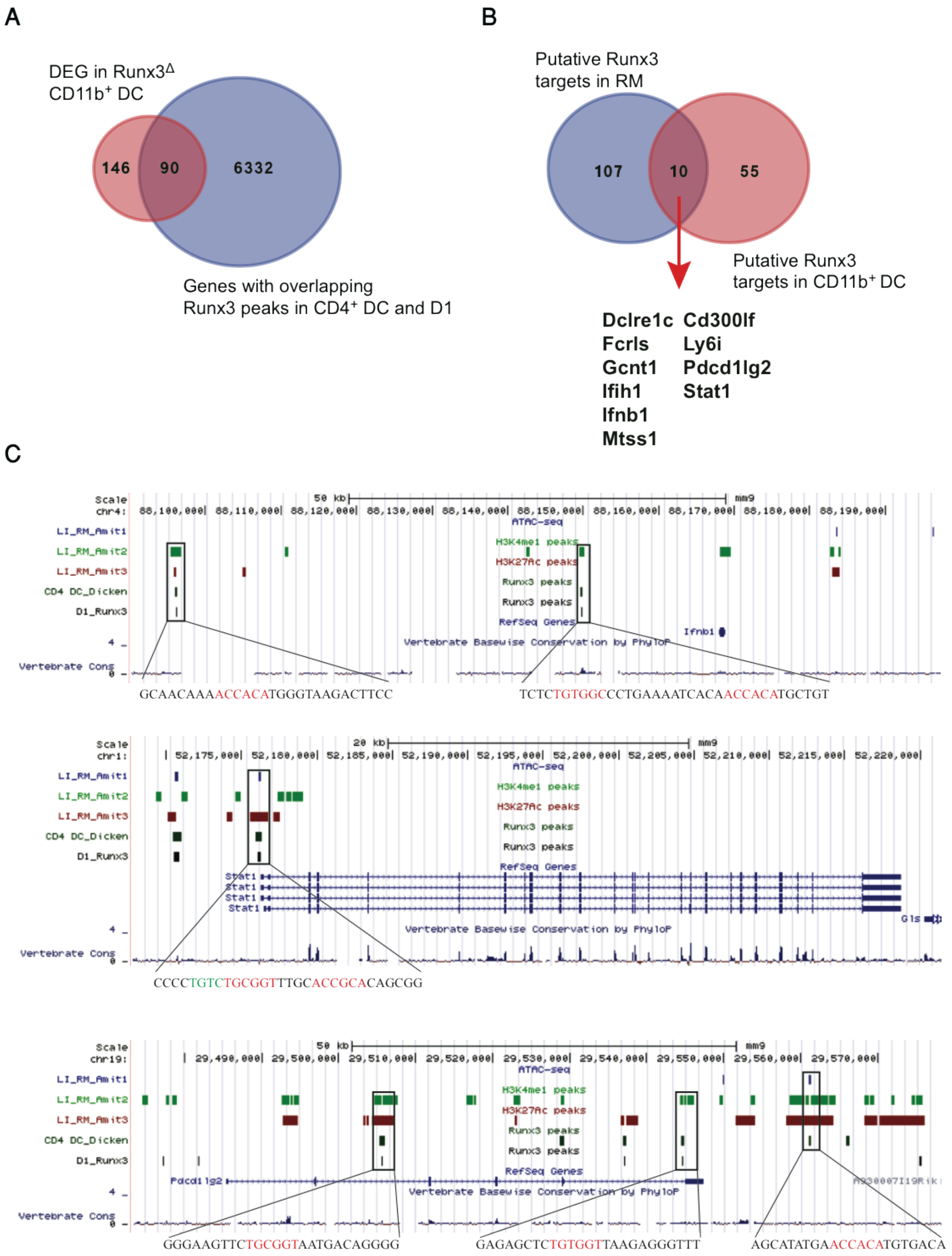

**Figure S6**. **Putative Runx3 target genes in colonic CD11b$^{+}$ DC**. **A,** Venn diagram depicting cross-analysis of DEGs in colonic Runx3$^{\Delta}$ CD11b$^{+}$ DC with genes harboring Runx3 peaks in D1 cells and splenic CD4$^{+}$ DC. **B,** Venn diagram depicting cross-analysis of Runx3 target genes in CD11b$^{+}$ DC and Runx3 target genes in colonic RM. **C,** D1 cells and splenic CD4$^{+}$ DC Runx3 occupied regions in three high-confidence DEGs common to colonic CD11b$^{+}$ DC and RM.

**Table S1-DEG RM**

**Runx3_cKO_Down regulated** **Runx3_cKO_Up regulated**

**(Fold-change >= 1.5, p-value <=0.05)**

| | | |
|---|---|---|
| 100043387 | Jmy | AA467197 |
| 0610010B08Rik | Kcnj16 | Acsl1 |
| 1300014I06Rik | Kmo | Adora2a |
| 2900026A02Rik | Leprel1 | Arg1 |
| 4632434I11Rik | Lpar6 | Arl5a |
| A630033H20Rik | Lphn3 | Arrdc4 |
| A630072M18Rik | Man1c1 | Ass1 |
| Abca9 | Map3k2 | Batf2 |
| Abhd15 | Mctp1 | Bcl2l14 |
| Acss1 | Med29 | Btnl5 |
| Adam22 | Mmp13 | C3 |
| Arhgap18 | Ms4a14 | Car2 |
| Arhgef3 | Mtss1 | Ccl17 |
| Arl4d | Mylip | Ccl22 |
| Asb2 | Naf1 | Ccl24 |
| Atxn1 | Nol8 | Cd300lf |
| B3galt5 | Nr1d2 | Cfb |
| BC055004 | Nt5c3 | Clca4 |
| Bend6 | Nt5dc2 | Clec12a |
| Bhlhe41 | Ntpcr | Cxcl9 |
| C6 | Ogn | Ddhd1 |
| Capn3 | P2rx1 | Dmbt1 |
| Ccl12 | Pde7b | Enc1 |
| Ccl3 | Pgf | F10 |
| Ccl4 | Plau | Flt1 |
| Cd59a | Pmepa1 | Gbp6 |
| Cfh | Ppp1r15a | Glipr2 |
| Ch25h | Rad51c | Gm12185 |
| Cited2 | Rcan1 | Gm4841 |
| Clec4b1 | Rcbtb2 | Gpnmb |
| Col14a1 | Reps2 | Gpx2 |
| Ctns | Sap30 | Gzma |
| Ctsf | Sass6 | Hif1a |
| Dcakd | Sepn1 | Iigp1 |
| Dck | Sh2d1b1 | Il12a |
| Dclre1c | Siglech | Il4i1 |
| Ddx60 | Slamf9 | Inhba |
| Dnajb4 | Slc25a23 | Insl6 |
| Dtx4 | Slc39a8 | Irg1 |
| Dusp18 | Slc40a1 | Layn |
| Emp1 | Slc9a3r2 | Ly6i |
| Extl2 | Spire1 | Ly75 |
| F630028O10Rik | Ssh2 | Marco |
| Fads3 | St3gal6 | Mgst1 |
| Fam125b | St5 | Mif |

| | | |
|---|---|---|
| Fam129a | Tanc2 | Nlrc5 |
| Fam189a2 | Tcp11l2 | Nos2 |
| Fap | Thap6 | Nufip1 |
| Fblim1 | Timd4 | Olfr566 |
| Fcrls | Tnf | Pdcd1lg2 |
| Fgf11 | Tnfrsf23 | Pfkp |
| Gcnt1 | Tnfsf4 | Phlpp1 |
| Gm6377 | Usp18 | Prdx5 |
| Gpr114 | Ust | Procr |
| Gpr34 | Xlr | Rffl |
| Gpx3 | Zbtb10 | Scarf1 |
| Grasp | Zfp758 | Sdc1 |
| Hnmt | Zfp931 | Sema4a |
| Hpgd | | Serpinb6b |
| Hpgds | | Serpinb9 |
| I830077J02Rik | | Slc7a11 |
| Ifih1 | | Slc7a2 |
| Ifit1 | | Stat1 |
| Ifit3 | | Tap1 |
| Ifnb1 | | Tarm1 |
| Il10 | | Tgtp1 |
| Il1r1 | | Tpi1 |
| Il22ra2 | | Tspan33 |
| Ints6 | | Upp1 |
| Ipcef1 | | Vdr |

**Table S1-putative Runx3 target genes in RM**

117 DEGs with overlapping peaks containg RUNX motif

| Gene name | SNP | Disease | SNP | Disease | SNP | Disease |
|---|---|---|---|---|---|---|
| 2900026A02Rik | | | | | | |
| AA467197 | | | | | | |
| Abhd15 | | | | | | |
| Acsl1 | | | | | | |
| Acss1 | | | | | | |
| Adam22 | | | | | | |
| Arg1 | | | | | | |
| Arhgap18 | | | | | | |
| Arhgef3 | | | | | | |
| Arl4d | | | | | | |
| Arl5a | | | | | | |
| Arrdc4 | | | | | | |
| Asb2 | | | | | | |
| Ass1 | | | | | | |
| Atxn1 | | | | | | |
| Batf2 | | | | | | |
| Bcl2l14 | | | | | | |
| Bend6 | | | | | | |
| C3 | | | | | | |
| C6 | | | | | | |
| Capn3 | | | | | | |
| Ccl12 | | | | | | |
| Ccl22 | | | | | | |
| Ccl3 | | | | | | |
| Ccl4 | | | | | | |
| Cd300lf | rs10512597 | CD | | | | |
| Cfb | rs4151657 | UC | | | | |
| Ch25h | | | | | | |
| Cited2 | | | | | | |
| Clec12a | | | | | | |
| Col14a1 | | | | | | |
| Ctns | | | | | | |
| Ctsf | | | | | | |
| Dcakd | | | | | | |
| Dck | | | | | | |
| Dclre1c | | | | | | |
| Ddhd1 | | | | | | |
| Dnajb4 | | | | | | |
| Dtx4 | | | | | | |
| Dusp18 | | | | | | |
| Emp1 | | | | | | |
| Enc1 | | | | | | |
| Extl2 | | | | | | |
| F630028O10Rik | | | | | | |
| Fam125b | | | | | | |
| Fam129a | | | | | | |

| | | | | | | |
|---|---|---|---|---|---|---|
| Fblim1 | | | | | | |
| Fcrls | | | | | | |
| Gcnt1 | | | | | | |
| Glipr2 | | | | | | |
| Gpr114 | | | | | | |
| Gpx3 | | | | | | |
| Grasp | | | | | | |
| Hpgd | | | | | | |
| Hpgds | | | | | | |
| I830077J02Rik | | | | | | |
| Ifih1 | rs2111485 | IBD | rs1990760 | IBD | rs503747517 | CD, UC |
| Ifnb1 | | | | | | |
| Il10 | rs30224505 | IBD, CD, UC | rs3024493 | IBD, UC | rs12075255 | CD, UC |
| Il22ra2 | | | | | | |
| Il4i1 | | | | | | |
| Inhba | | | | | | |
| Ipcef1 | | | | | | |
| Irg1 | | | | | | |
| Jmy | | | | | | |
| Kcnj16 | | | | | | |
| Kmo | | | | | | |
| Layn | | | | | | |
| Leprel1 | | | | | | |
| Lpar6 | | | | | | |
| Ly6i | | | | | | |
| Man1c1 | | | | | | |
| Map3k2 | | | | | | |
| Mctp1 | | | | | | |
| Med29 | | | | | | |
| Mmp13 | | | | | | |
| Ms4a14 | | | | | | |
| Mtss1 | | | | | | |
| Mylip | | | | | | |
| Naf1 | | | | | | |
| Nlrc5 | | | | | | |
| Nos2 | rs9797244 | CD, UC | rs28998802 | CD, UC | rs2779255 | CD, UC |
| Nr1d2 | | | | | | |
| Nt5c3 | | | | | | |
| Nt5dc2 | | | | | | |
| Ntpcr | | | | | | |
| Nufip1 | | | | | | |
| Pdcd1lg2 | | | | | | |
| Pde7b | | | | | | |
| Pfkp | | | | | | |
| Phlpp1 | | | | | | |
| Plau | rs2688608 | IBD | rs2227564 | IBD | rs2227551 | IBD, CD, UC |
| Pmepa1 | | | | | | |
| Prdx5 | rs694739 | CD | | | | |

| | | |
|---|---|---|
| Rcan1 | | |
| Rcbtb2 | | |
| Rffl | | |
| Sdc1 | | |
| Sema4a | | |
| Sepn1 | | |
| Serpinb6b | | |
| Serpinb9 | | |
| Sh2d1b1 | | |
| Siglech | | |
| Slamf9 | | |
| Slc40a1 | | |
| Slc7a11 | | |
| Spire1 | | |
| Ssh2 | | |
| St3gal6 | | |
| Stat1 | | |
| Tanc2 | | |
| Tcp11l2 | | |
| Tnf | rs1799964 | CD |
| Tspan33 | | |
| Ust | | |
| Zbtb10 | | |

### Table S1-P4 upTGFb1R-cKO down RM

53 Common elements in "P1-P4_up_RM" and "Tgfb_down_RM"

| | |
|---|---|
| 2900026A02Rik | Mvb12b |
| 8430419L09Rik | Mylip |
| Abcc3 | Ntpcr |
| Adam19 | Pdgfb |
| Arhgap22 | Pdia4 |
| Capn3 | Pmepa1 |
| Cd276 | Pxdc1 |
| Cd33 | Rnf180 |
| Cpd | Sash1 |
| Cx3cr1 | Sema4b |
| Cxcl16 | Slco3a1 |
| D8Ertd82e | Smad7 |
| Dtx4 | Soga1 |
| Enpp4 | Spint1 |
| Ermap | Stard13 |
| F11r | Stk38l |
| Fam189a2 | Ston2 |
| Fam214a | Tgfbr1 |
| Fam46c | Tlr12 |
| Fbxo32 | Tmem119 |
| Gna12 | Tnfsf13b |
| Hes1 | Vipr1 |
| Hic1 | Zfp691 |
| Itgav | Zmynd15 |
| Itgb5 | |
| Kcnj10 | |
| Kynu | |
| Mgat4a | |
| Mmp13 | |

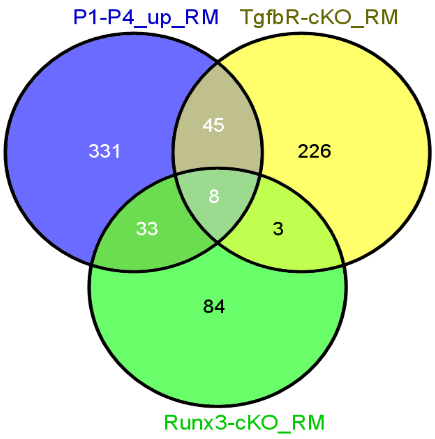

8 Common elements in "P1-P4_up_RM", "Tgfb_down_RM" and "Rx3_down RM":

2900026A02Rik
Capn3
Dtx4
Fam189a2
Mmp13
Mylip
Ntpcr
Pmepa1

### Table S1-DEG CD11b+ DC

**Runx3_cKO_Down regulated** **Runx3_cKO_Up regulated**

**(Fold-change >= 1.5, p-value <=0.05)**

| | | |
|---|---|---|
| Abca9 | Kitl | 2610528A11Rik |
| Acer2 | Leprel1 | 9330175E14Rik |
| Ago3 | Lifr | Abhd6 |
| Ahnak | Lilra6 | Aif1 |
| Ak8 | Lima1 | Ass1 |
| Akr1c18 | Lmna | Batf2 |
| Aldh1a2 | LOC100503338 | Bcl2l14 |
| Alox15 | Lphn2 | Bex6 |
| Alox5 | Lrrc1 | C3 |
| Ankrd12 | Ltb4r1 | Car2 |
| Anxa1 | Lyz1 | Ccl8 |
| Anxa2 | Mamdc2 | Cd300lf |
| Arhgap10 | Mamld1 | Cd3g |
| Arhgap21 | Man2a2 | Cdh17 |
| Arhgap26 | Mir18 | Ceacam16 |
| Arhgap39 | Mir221 | Cfb |
| Aspn | Ms4a14 | Col17a1 |
| Atp10d | Ms4a7 | Cox6a2 |
| Bco2 | Ms4a8a | Cxcl9 |
| C3ar1 | Mt2 | Dmbt1 |
| Calcrl | Mtmr10 | Dnase1l3 |
| Camk2g | Mtmr14 | Ermap |
| Capn2 | Mtss1 | Esam |
| Capn5 | Mx2 | Gatm |
| Ccl7 | Myo9a | Gatsl3 |
| Cd163 | Ndnf | Gbp2 |
| Cd2 | Notch1 | Gca |
| Cd200r1 | Nsun6 | Gm14005 |
| Cd209b | Oasl1 | Gm16379 |
| Cd209c | Pdlim1 | Gm4951 |
| Cd226 | Plscr2 | Gpnmb |
| Cd24a | Plxna4 | Gpr33 |
| Cd300lb | Pmp22 | Gpx2 |
| Cd93 | Polg2 | Gstt1 |
| Cebpd | Prelid2 | Gzmb |
| Cgnl1 | Pros1 | Hap1 |
| Ciart | Ptger2 | Hbegf |
| Cks2 | Ptgir | Ifi47 |
| Crim1 | Rcan1 | Igkv4-59 |
| Dcbld2 | Retnla | Igkv4-69 |
| Dclre1c | Rnd3 | Igkv4-70 |
| Dgkg | Rps6ka2 | Igtp |
| Dhx40 | Satb2 | Il12a |
| Dlc1 | Sema6b | Itgad |
| Dpep2 | Sept11 | Ly6a |

| | | |
|---|---|---|
| Dusp18 | Serpinb1b | Ly6c1 |
| Ecm1 | Serpinb2 | Ly6c2 |
| Ednrb | Serpinb8 | Ly6i |
| Efnb2 | Sgms2 | Mcpt2 |
| Enkur | Slc13a3 | Mif |
| Ereg | Slc22a21 | Mpzl2 |
| F13a1 | Slc9a4 | Nkx2-9 |
| F2rl2 | Snord37 | Nlrc5 |
| F5 | Snord58b | Olfr521 |
| F630028O10Rik | Spink2 | Papss2 |
| Fam102b | Sult2b1 | Pdcd1lg2 |
| Fam20a | Syne2 | Pigr |
| Fbxo30 | Tespa1 | Pla2g16 |
| Fcrls | Tmem167 | Pla2g2d |
| Fgfr1 | Tnfsf14 | Pla2g4e |
| Fn1 | Tppp3 | Plac8 |
| Furin | Ttc39c | Plcb4 |
| Gabbr1 | Unc119b | Plekhs1 |
| Gch1 | Vav3 | Prrg4 |
| Gcnt1 | Vcl | Ptger3 |
| Gm10099 | Vmn2r96 | Rragd |
| Gm10752 | Xlr | Serpina3f |
| Gm25323 | Zbtb32 | Slc27a2 |
| Gm9883 | | Slc28a3 |
| Gpr116 | | Slc4a8 |
| Gpr56 | | Snap47 |
| Hr | | Sprr1a |
| Ifih1 | | Spsb1 |
| Ifnb1 | | Src |
| Igkv10-96 | | St14 |
| Il1r1 | | Stap2 |
| Il1rap | | Stat1 |
| Il1rl1 | | Tbx21 |
| Il3 | | Tgm2 |
| Il33 | | Tgtp2 |
| Il6 | | Tm4sf5 |
| Irf4 | | Tvp23a |
| Itpkb | | Ube2l6 |
| Jam2 | | Vcam1 |

#### Table S1-Putative Runx3 targets DC

##### 65 DEGs with overlapping peaks containing RUNX motif

| Gene name | SNP | Disease | SNP | Disease | SNP | Disease |
|---|---|---|---|---|---|---|
| Ahnak | | | | | | |
| Alox5 | | | | | | |
| Anxa2 | | | | | | |
| Arhgap10 | | | | | | |
| Arhgap26 | | | | | | |
| Arhgap39 | | | | | | |
| Bco2 | | | | | | |
| Bex6 | | | | | | |
| C3ar1 | | | | | | |
| Camk2g | | | | | | |
| Cd300lb | | | | | | |
| Cd300lf | rs10512597 | CD | | | | |
| Ceacam16 | | | | | | |
| Cgnl1 | | | | | | |
| Cks2 | | | | | | |
| Crim1 | | | | | | |
| Dclre1c | | | | | | |
| Dgkg | | | | | | |
| Dhx40 | | | | | | |
| Ecm1 | | | | | | |
| F13a1 | | | | | | |
| F2rl2 | | | | | | |
| Fam102b | | | | | | |
| Fcrls | | | | | | |
| Fgfr1 | | | | | | |
| Furin | | | | | | |
| Gca | | | | | | |
| Gcnt1 | | | | | | |
| Gpr56 | | | | | | |
| Hap1 | | | | | | |
| Hbegf | | | | | | |
| Ifi47 | | | | | | |
| Ifih1 | rs2111485 | IBD | rs1990760 | IBD | rs503747517 | CD, UC |
| Ifnb1 | | | | | | |
| Il1rap | | | | | | |
| Irf4 | rs7773324 | IBD, CD | rs1033180 | Celiac | rs1050976 | Celiac |
| Itgad | | | | | | |
| Jam2 | | | | | | |
| Lmna | | | | | | |
| Ly6a | | | | | | |
| Ly6i | | | | | | |
| Ms4a7 | | | | | | |
| Mtmr14 | | | | | | |

| | | | | |
|---|---|---|---|---|
| Mtss1 | | | | |
| Pdcd1lg2 | | | | |
| Plac8 | | | | |
| Plxna4 | | | | |
| Polg2 | | | | |
| Prelid2 | | | | |
| Ptger3 | | | | |
| Sema6b | | | | |
| Serpinb8 | | | | |
| Slc22a21 | rs6596075 | CD | rs1016988 | CD |
| Slc27a2 | | | | |
| Slc28a3 | | | | |
| Snap47 | | | | |
| Src | | | | |
| Stat1 | | | | |
| Tgm2 | | | | |
| Tm4sf5 | | | | |
| Tmem167 | | | | |
| Tvp23a | | | | |
| Vav3 | | | | |
| Vcl | | | | |
| Vmn2r96 | | | | |

### Table S1-Panther pathways

H3K27ac marked peaks_intestinal RM

**PANTHER Pathway (20+ terms)** Global controls

*Table controls:* Export | Shown top rows in this table: 20 Set | Term annotation count: Min: 1 Max: Inf Set | Visualize this table: [select one]

| Term Name | Binom Rank | Binom Raw P-Value | *Binom FDR Q-Val* | Binom Fold Enrichment | Binom Observed Region Hits | Binom Region Set Coverage | Hyper Rank | *Hyper FDR Q-Val* | Hyper Fold Enrichment | Hyper Observed Gene Hits | Hyper Total Genes | Hyper Gene Set Coverage |
|---|---|---|---|---|---|---|---|---|---|---|---|---|
| Inflammation mediated by chemokine and cytokine signaling pathway | 1 | 3.5540e-183 | 5.3666e-181 | 2.7493 | 1,090 | 3.25% | 12 | 1.3857e-4 | 1.2251 | 163 | 217 | 1.26% |
| Apoptosis signaling pathway | 2 | 6.8467e-141 | 5.1692e-139 | 3.3071 | 634 | 1.89% | 2 | 2.4676e-11 | 1.4975 | 101 | 110 | 0.78% |
| B cell activation | 3 | 7.0519e-134 | 3.5494e-132 | 4.0370 | 468 | 1.40% | 5 | 7.5537e-8 | 1.5480 | 56 | 59 | 0.43% |
| T cell activation | 4 | 2.0424e-100 | 7.7100e-99 | 3.1567 | 481 | 1.43% | 3 | 1.1643e-9 | 1.5439 | 71 | 75 | 0.55% |
| PDGF signaling pathway | 5 | 4.3286e-96 | 1.3072e-94 | 2.4840 | 683 | 2.04% | 1 | 1.1237e-11 | 1.4851 | 112 | 123 | 0.86% |
| EGF receptor signaling pathway | 8 | 1.3712e-77 | 2.5882e-76 | 2.4159 | 580 | 1.73% | 4 | 7.9995e-8 | 1.4215 | 95 | 109 | 0.73% |
| Ras Pathway | 9 | 4.6481e-76 | 7.7985e-75 | 2.9614 | 399 | 1.19% | 6 | 1.5499e-7 | 1.5092 | 62 | 67 | 0.48% |
| Interleukin signaling pathway | 10 | 2.6386e-72 | 3.9842e-71 | 2.7272 | 433 | 1.29% | 13 | 2.3139e-4 | 1.3377 | 73 | 89 | 0.56% |
| Angiogenesis | 11 | 6.9372e-70 | 9.5229e-69 | 2.0470 | 749 | 2.23% | 15 | 7.9869e-4 | 1.2391 | 117 | 154 | 0.90% |
| Toll receptor signaling pathway | 12 | 7.0478e-65 | 8.8685e-64 | 3.4905 | 268 | 0.80% | 7 | 1.1064e-6 | 1.5912 | 40 | 41 | 0.31% |
| p38 MAPK pathway | 13 | 3.7626e-63 | 4.3704e-62 | 3.5711 | 253 | 0.75% | 10 | 9.2011e-5 | 1.5403 | 34 | 36 | 0.26% |
| VEGF signaling pathway | 14 | 3.1940e-52 | 3.4450e-51 | 2.6626 | 323 | 0.96% | 28 | 4.1467e-3 | 1.3269 | 48 | 59 | 0.37% |
| Huntington disease | 15 | 3.9907e-51 | 4.0173e-50 | 2.1351 | 492 | 1.47% | 22 | 1.7487e-3 | 1.2504 | 92 | 120 | 0.71% |
| FGF signaling pathway | 16 | 1.7067e-47 | 1.6107e-46 | 2.0608 | 496 | 1.48% | 16 | 8.0179e-4 | 1.2836 | 85 | 108 | 0.65% |
| Oxidative stress response | 17 | 9.8267e-47 | 8.7285e-46 | 2.7958 | 265 | 0.79% | 9 | 2.4751e-5 | 1.4978 | 45 | 49 | 0.35% |
| Parkinson disease | 18 | 1.5153e-46 | 1.2711e-45 | 2.3685 | 357 | 1.06% | 27 | 2.6509e-3 | 1.2788 | 69 | 88 | 0.53% |
| Interferon-gamma signaling pathway | 19 | 1.1818e-31 | 9.3923e-31 | 2.8468 | 171 | 0.51% | 32 | 1.1041e-2 | 1.4135 | 26 | 30 | 0.20% |
| Angiotensin II-stimulated signaling through G proteins and beta-arrestin | 22 | 1.5834e-27 | 1.0868e-26 | 2.6532 | 166 | 0.49% | 37 | 3.4322e-2 | 1.3224 | 30 | 37 | 0.23% |
| p53 pathway feedback loops 2 | 24 | 2.0604e-24 | 1.2963e-23 | 2.2108 | 209 | 0.62% | 11 | 1.0261e-4 | 1.4860 | 41 | 45 | 0.32% |
| JAK/STAT signaling pathway | 26 | 1.9199e-22 | 1.1150e-21 | 3.5155 | 87 | 0.26% | 25 | 2.4017e-3 | 1.6309 | 16 | 16 | 0.12% |

Runx3-bound overlapping peaks_CD4+ DC and D1

**PANTHER Pathway (20+ terms)** Global controls

*Table controls:* Export | Shown top rows in this table: 20 Set | Term annotation count: Min: 1 Max: Inf Set | Visualize this table: [select one]

| Term Name | Binom Rank | Binom Raw P-Value | *Binom FDR Q-Val* | Binom Fold Enrichment | Binom Observed Region Hits | Binom Region Set Coverage | Hyper Rank | *Hyper FDR Q-Val* | Hyper Fold Enrichment | Hyper Observed Gene Hits | Hyper Total Genes | Hyper Gene Set Coverage |
|---|---|---|---|---|---|---|---|---|---|---|---|---|
| Inflammation mediated by chemokine and cytokine signaling pathway | 1 | 6.3308e-32 | 9.5595e-30 | 2.5493 | 206 | 3.01% | 3 | 3.4229e-8 | 1.6527 | 109 | 217 | 1.69% |
| T cell activation | 2 | 1.3533e-25 | 1.0217e-23 | 3.3808 | 105 | 1.54% | 1 | 6.1465e-10 | 2.2812 | 52 | 75 | 0.81% |
| Apoptosis signaling pathway | 3 | 1.5574e-25 | 7.8387e-24 | 3.0710 | 120 | 1.76% | 7 | 6.6959e-6 | 1.7648 | 59 | 110 | 0.92% |
| B cell activation | 4 | 8.1535e-24 | 3.0779e-22 | 3.6820 | 87 | 1.27% | 6 | 1.7846e-6 | 2.1191 | 38 | 59 | 0.59% |
| Gonadotropin-releasing hormone receptor pathway | 5 | 3.4511e-22 | 1.0422e-20 | 2.0108 | 232 | 3.39% | 2 | 1.5244e-9 | 1.6877 | 119 | 232 | 1.85% |
| Angiogenesis | 6 | 1.7686e-18 | 4.4509e-17 | 2.1588 | 161 | 2.36% | 5 | 5.4522e-7 | 1.7092 | 80 | 154 | 1.24% |
| PDGF signaling pathway | 7 | 2.9384e-18 | 6.3385e-17 | 2.3553 | 132 | 1.93% | 4 | 1.0135e-7 | 1.8457 | 69 | 123 | 1.07% |
| EGF receptor signaling pathway | 8 | 3.4280e-14 | 6.4703e-13 | 2.2480 | 110 | 1.61% | 9 | 6.1883e-5 | 1.6904 | 56 | 109 | 0.87% |
| VEGF signaling pathway | 10 | 1.5088e-12 | 2.2784e-11 | 2.7097 | 67 | 0.98% | 12 | 4.9795e-4 | 1.8403 | 33 | 59 | 0.51% |
| Toll receptor signaling pathway | 11 | 1.1925e-11 | 1.6370e-10 | 3.1311 | 49 | 0.72% | 17 | 4.8352e-3 | 1.8457 | 23 | 41 | 0.36% |
| Ras Pathway | 12 | 1.8701e-11 | 2.3532e-10 | 2.5126 | 69 | 1.01% | 10 | 3.0531e-4 | 1.8170 | 37 | 67 | 0.57% |
| Huntington disease | 13 | 4.5326e-11 | 5.2648e-10 | 2.0866 | 98 | 1.43% | 14 | 2.7569e-3 | 1.5080 | 55 | 120 | 0.85% |
| Oxidative stress response | 14 | 6.8882e-11 | 7.4294e-10 | 2.7951 | 54 | 0.79% | 23 | 1.3107e-2 | 1.6787 | 25 | 49 | 0.39% |
| PI3 kinase pathway | 15 | 7.0413e-11 | 7.0882e-10 | 2.7625 | 55 | 0.80% | 26 | 4.6380e-2 | 1.5766 | 23 | 48 | 0.36% |
| p38 MAPK pathway | 17 | 8.9991e-10 | 7.9933e-9 | 2.9778 | 43 | 0.63% | 11 | 4.6314e-4 | 2.1021 | 23 | 36 | 0.36% |
| Interleukin signaling pathway | 18 | 2.7278e-9 | 2.2883e-8 | 2.1940 | 71 | 1.04% | 15 | 2.8912e-3 | 1.5897 | 43 | 89 | 0.67% |
| Interferon-gamma signaling pathway | 19 | 2.7859e-9 | 2.2141e-8 | 3.1038 | 38 | 0.56% | 19 | 5.8666e-3 | 1.9741 | 18 | 30 | 0.28% |
| Histamine H1 receptor mediated signaling pathway | 21 | 1.3750e-8 | 9.8866e-8 | 2.4867 | 50 | 0.73% | 24 | 1.4958e-2 | 1.7235 | 22 | 42 | 0.34% |
| Axon guidance mediated by semaphorins | 22 | 1.4854e-8 | 1.0195e-7 | 3.6126 | 28 | 0.41% | 13 | 1.3835e-3 | 2.5161 | 13 | 17 | 0.20% |
| Cytoskeletal regulation by Rho GTPase | 23 | 3.1556e-8 | 2.0717e-7 | 2.3495 | 53 | 0.78% | 25 | 1.6875e-2 | 1.5483 | 32 | 68 | 0.50% |
